# The X-factor in ART: does the use of Assisted Reproductive Technologies influence DNA methylation on the X chromosome?

**DOI:** 10.1101/2022.10.06.510603

**Authors:** Julia Romanowska, Haakon E. Nustad, Christian M. Page, William R.P. Denault, Jon Bohlin, Yunsung Lee, Maria C. Magnus, Kristine L. Haftorn, Miriam Gjerdevik, Boris Novakovic, Richard Saffery, Håkon K. Gjessing, Robert Lyle, Per Magnus, Siri E. Håberg, Astanand Jugessur

**Author notes:** Corresponding author:* Julia Romanowska, PhD, Department of Global Public Health and Primary Care, University of Bergen, 5020 Bergen, Norway.

## Abstract

**Background:** Assisted reproductive technologies (ART) may perturb DNA methylation (DNAm) in early embryonic development. Although a handful of epigenome-wide association studies of ART have been published, none have investigated CpGs on the X chromosome. To bridge this knowledge gap, we leveraged one of the largest collections of mother-father-newborn trios of ART and non-ART (natural) conceptions to date to investigate DNAm differences on the X chromosome.

**Materials and Methods:** The discovery cohort consisted of 982 ART and 963 non-ART trios from the Norwegian Mother, Father, and Child Cohort Study (MoBa). The replication cohort consisted of 149 ART and 58 non-ART neonates from the Australian “Clinical review of the Health of adults conceived following Assisted Reproductive Technologies” (CHART) study. The Illumina EPIC array was used to measure DNA methylation (DNAm) in both datasets. In the MoBa cohort, we performed a set of X-chromosome-wide association studies (“XWASs” hereafter) to search for sex-specific DNAm differences between ART and non-ART newborns. We tested several models to investigate the influence of various confounders, including parental DNAm. We also searched for differentially methylated regions (DMRs) and regions of co-methylation flanking the most significant CpGs. For replication purposes, we ran an analogous model to our main model on the CHART dataset.

**Results and conclusions:** In the MoBa cohort, we found more differentially methylated CpGs and DMRs in girls than boys. Most of the associations persisted even after controlling for parental DNAm and other confounders. Many of the significant CpGs and DMRs were in gene-promoter regions, and several of the genes linked to these CpGs are expressed in tissues relevant for both ART and sex (testis, placenta, and fallopian tube). We found no support for parental infertility as an explanation for the observed associations in the newborns. The most significant CpG in the boys-only analysis was in *UBE2DNL*, which is expressed in testes but with unknown function. The most significant CpGs in the girls-only analysis were in *EIF2S3* and *AMOT*. These three loci also displayed differential DNAm in the CHART cohort. Overall, genes that co-localized with the significant CpGs and DMRs are implicated in several key biological processes (e.g., neurodevelopment) and disorders (e.g., intellectual disability and autism. These connections are particularly compelling in light of previous findings indicating that neurodevelopmental outcomes differ in ART-conceived children compared to naturally-conceived.

## 1 Background

The use of assisted reproductive technologies (ART) has been on the rise in most parts of the world since the first baby was born to *in vitro* fertilization (IVF) in 1978^1;2^. The trend of declining fecundity and greater reliance on ART to conceive is expected to persist in the future, as egg-freezing gains more acceptance in contemporary societies and more couples choose to postpone childbearing^3–5^. As the clinical and laboratory procedures for ART coincide with the developmental window in which the early embryo undergoes extensive epigenetic remodeling^6–8^, it is critical to determine whether the ART procedures themselves or some underlying mechanisms related to parental characteristics (e.g., parental infertility) are responsible for the observed epigenetic differences between ART and non-ART newborns. A number of epigenome-wide association studies (EWASs) of ART have been published in recent years^9–17^ and have already contributed substantially to our current understanding of epigenetic changes associated with ART. However, none of these studies have investigated the effect of epigenetic markers on the X chromosome.

Until recently, most genome-wide association studies (GWASs) were also performed almost exclusively on autosomes, leaving out single-nucleotide polymorphisms (SNPs) on the X chromosome, even though this chromosome constitutes ~5% of the human genome and houses ~1000 genes, several of which have been associated with complex traits^18;19^. The main reason for this exclusion is that the initial methods for GWAS were primarily designed for autosomal markers, as analyzing different X chromosome contents in males and females comes with its own set of analytic challenges^20^. To fill this knowledge gap, we and others have developed a suite of biostatistical tools for analyzing X-linked SNPs both individually and as haplotypes^21–31^. Currently, there is a similar trend of systematic exclusion of CpGs on the sex chromosomes in the vast majority of EWASs, which may result in overlooking important associations.

There are several reasons why X chromosome markers are less tractable to analyze than autosomal markers. First, one needs to account for X chromosome inactivation (XCI) in which one of the X chromosomes in female somatic cells is randomly selected and transcriptionally inactivated in early embryonic development^32;33^. This crucial mechanism ensures a balanced dosage of X-linked genes in males and females^34–36^. However, XCI is not complete in humans, with approximately 12% of the genes reported to escape XCI and a further 15% differing in their XCI status across individuals, tissues, and cells^33;37–39^. Second, the analysis of X-linked markers is complicated by genes in the pseudoautosomal regions (PARs) which are expressed in a similar fashion to autosomal genes as a consequence of escaping XCI^40;41^. Third, the gradual loss of the X chromosome with age^42^ may further complicate the analysis of X-linked markers when comparing cohorts that differ significantly with age.

Despite these challenges, taking X chromosome markers into account in a GWAS or EWAS is important based on the following observations: (a) genes on the X chromosome are known to play essential roles in transcriptional regulation of autosomal genes^43;44^, (b) several traits show a consistently higher prevalence in one sex, and (c) there are distinct physical differences between the sexes (sexual dimorphism)^45^. All of these features might stem from sex-specific differences and this is especially relevant for differences occurring *prior* to gonadal differentiation, i.e., differences that are solely attributable to sex chromosome content rather than those induced by gonadal and hormonal changes^34;35;46^. Although a wide variety of traits are known to exhibit sex-specific DNA methylation (DNAm) signatures on the autosomes^47–57^, less is known about the presence of such signatures on the X chromosome, possibly due to the overall lack of focus on X-linked markers and the dearth of X-chromosome-wide association studies (XWASs) conducted to date. The few XWASs published thus far include an investigation of CpGs influenced by cigarette smoking, an exploration of differential chronological aging in males versus females, and a study of DNAm changes associated with aging on the X and Y chromosomes^58–61^.

Given these important knowledge gaps, our main objective was to examine sex-specific differences in DNAm profiles on the X chromosome by contrasting ART and naturally-conceived newborns. We used one of the largest case-control collection of mother-father-newborn trios of ART and non-ART conceptions to date^62^, stemming from the Norwegian Mother, Father, and Child Cohort Study (MoBa)^63^. The analyses were stratified by sex and adjusted for potential confounding factors (mother’s age, smoking status, BMI, and primiparity, as well as parental DNAm at each CpG). For replication purposes, we analyzed data from the Australian ‘Clinical review of the Health of 22–33 years old conceived with and without ART’ (CHART) cohort^64;65^.

## 2 Results

In the discovery cohort (MoBa), we analyzed DNAm data from 982 ART and 963 non-ART mother-father-newborn trios. These data were generated on the Illumina EPIC platform using DNA extracted from peripheral blood in adults and cord blood in newborns (for details, see Methods and ref.^62^). Our main aim was to identify differences in DNAm in ART versus non-ART newborns, both at a single-CpG level across the entire X chromosome (XWAS) and at a regional level where we searched for differentially methylated regions (DMRs). All the analyses were performed separately for boys and girls. In the main model, we adjusted for known confounders (mother’s age, smoking status, BMI, and primiparity). In addition, we tested three adjusted models, where we included: *(i)* parental DNAm at each CpG, *(ii)* birthweight and gestational age of the newborn, and *(iii)* all three covariates from *(i)* and *(ii)*. All the analyses were stratified by sex. We also explored co-methylation patterns between significant CpGs identified by the above analyses as well as other CpGs in the immediate flanking regions.

In the replication cohort (CHART), we analyzed DNAm data, also generated on the EPIC array, from 149 ART-conceived and 58 non-ART newborns. These analyses are outlined in Figure 1 and detailed in Materials and Methods.

**Figure 1:**
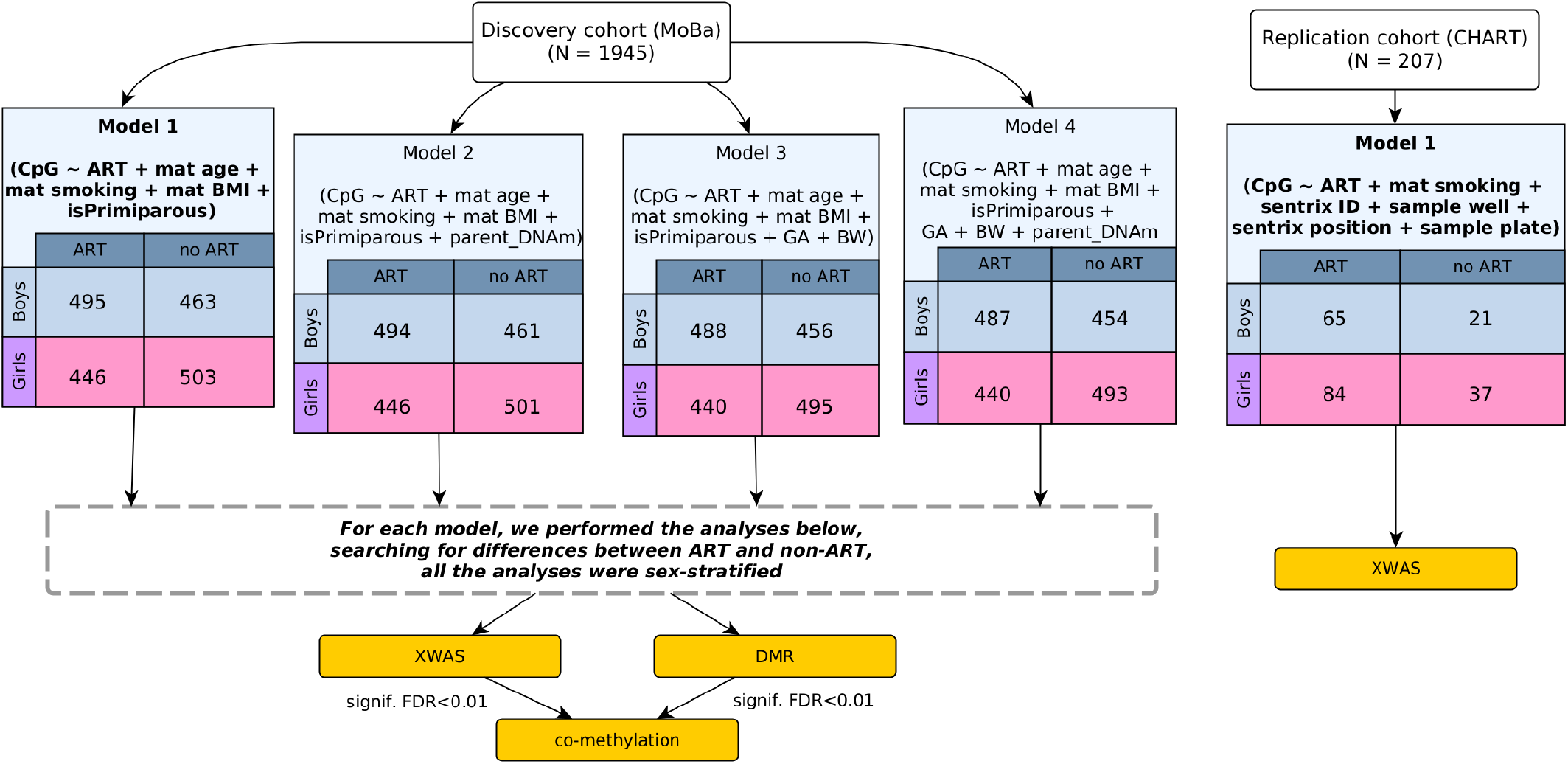
Overview of the analytic pipeline. We refer to **Model 1** as the ‘main model’ throughout this paper and focus primarily on significant findings from this model. The other models are referred to as ‘adjusted models’ and treated as sensitivity analyses. Abbreviations used in the figure: CpG = cytosine-phosphate-guanine; ART = assisted reproductive technologies; mat = maternal; BMI = body mass index; BW = birthweight; GA = gestational age; XWAS = X-chromosome-wide association study; DMR = differentially methylated region; FDR = false discovery rate; MoBa = The Mother, Father, and Child Cohort Study; CHART = The “Clinical review of the Health of adults conceived following Assisted Reproductive Technologies” study.

### 2.1 Differences between ART and non-ART newborns and their parents in the MoBa cohort

The ART parents were older than the non-ART parents, and the ART newborns weighed less than the non-ART newborns (Table 1). Fewer of the ART mothers smoked during pregnancy than the non-ART mothers, but, intriguingly, a higher proportion of the ART mothers were *past* smokers.

**Table 1:** Characteristics of the discovery cohort (MoBa).

| Characteristics | Non-ART (N = 983)<sup>1</sup> | ART (N = 962)<sup>1</sup> | p-value<sup>2</sup> |
| --- | --- | --- | --- |
| Male (%) | 470 (48%) | 505 (52%) | 0.039 |
| Maternal age (years) | 30 (27-33) | 33 (31-36) | <0.001 |
| Paternal age (years) | 32 (29-36) | 35 (32-38) | <0.001 |
| Gestational age (days) | 40 (39-41) | 40 (39-41) | 0.090 |
| (No data) | 4 | 0 |  |
| Birth weight (g) | 3,650 (3,330-3,970) | 3,540 (3,190-3,850) | <0.001 |
| (No data) | 0 | 1 |  |
| First child | 522 (53%) | 289 (30%) | <0.001 |
| Maternal BMI (kg/m<sup>2</sup>) | 23 (21-26) | 23 (21-26) | 0.4 |
| (No data) | 14 | 17 |  |
| Maternal smoking |  |  | <0.001 |
| Never | 490 (50%) | 494 (52%) |  |
| Past smoker | 253 (26%) | 358 (37%) |  |
| 1st trimester | 132 (13%) | 62 (6.5%) |  |
| 1st trimester and after | 104 (11%) | 44 (4.6%) |  |
| (No data) | 4 | 4 |  |

<sup>1</sup> n (%); median (IQR)
<sup>2</sup> Pearson's Chi-squared test; Wilcoxon rank-sum test

Figure 2 highlights the general trends in the XWAS results for boys and girls separately, before and after controlling for inflation using the R-package BACON (see Methods for details). Effect sizes for CpGs in the boys-only analyses showed a slight global hypermethylation (i.e., an overall higher DNAm level in the ART newborns). By contrast, the effect sizes in the girls-only analyses were dominated by global hypomethylation (overall lower DNAm level in the ART newborns). Figure 2 also illustrates the efficacy of BACON in reducing inflation in the p-values. A similar figure showing the results of all the models tested can be found in the Supplementary Figure S3.

**Figure 2:**
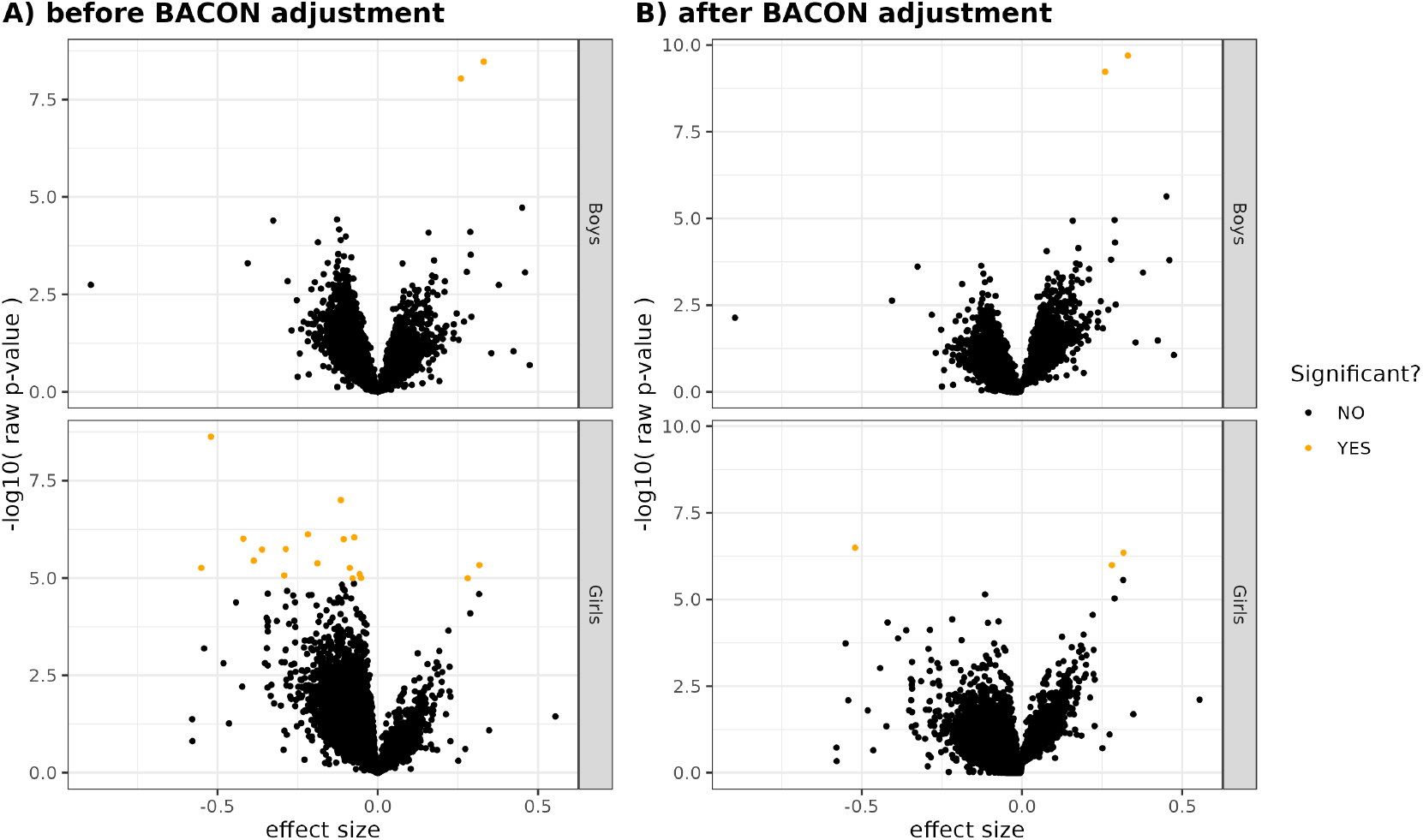
Effect sizes versus − log_10_ *p*-values for each of the X-linked CpGs included in the analyses of the MoBa cohort. Significant findings at FDR < 0.01 are highlighted in orange. The data are presented before (panel A) and after (panel B) p-value adjustment with the BACON algorithm.

### 2.2 Several CpGs were significantly differentially methylated between ART and non-ART in the MoBa cohort

We identified three significantly differentially methylated CpGs in the girls-only analysis (cg25034591, cg13866977, and cg26175661) and two CpGs in the boys-only analysis (cg00920314, cg04516011), all at a false discovery rate-adjusted (FDR-adjusted) p-value < 0.01. Strikingly, there was no overlap in the location of the significant findings between the girls-only and boys-only analyses (Figure 3 and Supplementary Figures S4-S6). A detailed summary of the significant results for all four statistical models is provided in the Supplementary Document 1. Additionally, tables with all the results are available in Github at https://github.com/folkehelseinstituttet/X-factor-ART.

**Figure 3:**
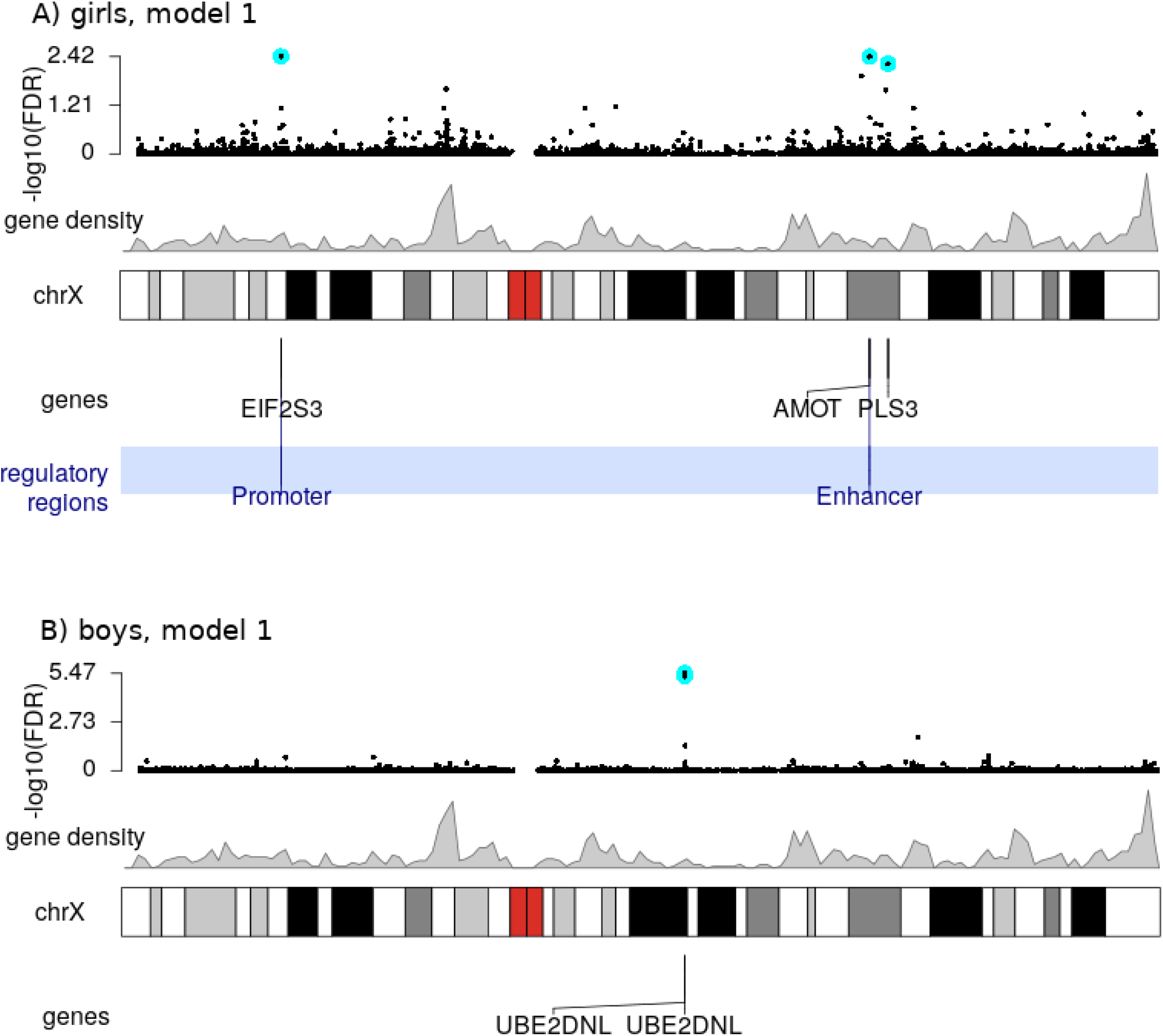
Results of the XWAS of girls (panel A) and boys (panel B) based on the main model (**Model 1**; `CpG` ~ `ART + maternal age + maternal smoking + maternal BMI + primiparity`). The top plot in each panel is a Manhattan plot of all the tested CpGs. The genomic locations of the most significant findings (FDR < 0.01) are highlighted by cyan-colored circles. Immediately beneath is a line plot of gene density, the chromosomal bands, and any genes and/or regulatory regions that overlap with the significant findings.

Adjusting for parental DNAm in the MoBa sample enabled ruling out parental characteristics as the reason for the observed DNAm differences. When the results of the main model were contrasted with those of the adjusted model, there was no significant change in the findings in the boys-only analyses. By contrast, only cg25034591 and cg13866977 remained significant across all models in the girls-only analyses. These results suggest that the differential methylation at these sites is more likely the result of the ART procedures themselves rather than parental DNAm.

We performed bootstrap analyses to evaluate the consistency with which the significant CpGs were retained. The two significant CpGs in the boys-only analyses (cg00920314 and cg04516011) showed a high degree of consistency. They were significant 54% and 47% of the time, respectively, which is substantially higher than the next CpG on the ranked list (cg00243584 at 9%). In the girls-only analyses, cg25034591 was significant in 51% of the bootstrap samples, but the other two CpGs were not as consistent (cg13866977 at 25% and cg26175661 at 19%, occupying positions six and 14 on the ranked list, respectively). The full list of CpGs found to be significant at least once, and the proportion of times a given CpG was found to be significant, are provided in Supplementary Data 1 for the boys-only analyses and Supplementary Data 2 for the girls-only analyses. These results are also provided in the Github repository.

The two significant CpGs detected in the boys-only analysis are adjacent and located within the gene ‘Ubiquitin conjugating enzyme E2 D N-terminal like’ (*UBE2DNL*) (Figure 3). In contrast, the significant CpGs in the girls-only analysis are located in different chromosomal regions, i.e., within ‘Eukaryotic translation initiation factor 2 subunit gamma’ (*EIF2S3*), ‘Ribosomal protein L18a pseudogene 15’ (*RPL18AP15*), ‘Angiomotin’ (*AMOT*), and ‘Plastin 3’ (*PLS3*) (Figure 3). Two of the CpGs are located within promoter regions and one within an enhancer. See Table 2 for a summary of the genes.

**Table 2:** Summary of the genes and loci identified in the current XWAS.

| Sex | Gene name (location; ensembl ID) | Full gene name (MIM entry) | Gene function<sup>1</sup> | Refs |
| --- | --- | --- | --- | --- |
| Girls | *EIF2S3*<br>(Xp22.11;<br>ENSG00000130741) | Eukaryotic translation initiation factor 2 subunit gamma (MIM:300161) | This gene encodes the core subunit of eukaryotic translation initiation factor-2 (eIF2). This gamma subunit is the largest component of a heterotrimeric GTP-binding protein which is essential for protein synthesis. Hemizygous mutations in *EIF2S3* cause an X-linked syndrome called 'mental retardation, epileptic seizures, hypogonadism and -genitalism, microcephaly, and obesity' (MEHMO). *EIF2S3* has also been reported to escape XCI. | 33;76;77<br>78;142;143 |
| Girls | *RPL18AP15*<br>(Xq23;<br>ENSG00000236189) | Ribosomal protein L18a pseudogene 15 | *RPL18AP15* is a pseudogene. | Not applicable |
| Girls | *AMOT*<br>(Xq23;<br>ENSG00000126016) | Angiomotin (MIM:300410) | This gene belongs to the motin family of angiostatin-binding proteins containing conserved coiled-coil domains and C-terminal PDZ binding motifs. *AMOT* is predominantly expressed in endothelial cells of capillaries and in larger vessels of the placenta where it may mediate the inhibitory effect of angiostatin on tube formation and the migration of endothelial cells toward growth factors during the formation of new blood vessels. | 144–146 |
| Girls | *PLS3*<br>(Xq23;<br>ENSG00000102024) | Plastin 3 (MIM:300131) | Plastins comprise a family of actin-binding proteins that are conserved throughout eukaryote evolution. They are expressed in most tissues of higher eukaryotes. Two ubiquitous plastin isoforms (L and T) have been identified in humans. |  |
| Boys | *UBE2DNL*<br>(Xq21.1;<br>ENSG00000229547) | Ubiquitin conjugating enzyme E2 D N-terminal like (pseudogene) | *UBE2DNL* is labeled 'pseudogene' in various gene databases, but it is reported to be expressed in testis. | Not applicable |

<sup>1</sup> Information on gene function was collated from various sources, including NCBI's Entrez Gene (https://www.ncbi.nlm.nih.gov/gene), Gene Cards (https://www.genecards.org/), the Online Inheritance in Man (OMIM) (https://omim.org/), and the cited references in the last column.

### 2.3 Patterns of co-methylation around the significant CpGs in the MoBa cohort

Analyzing clusters of DNAm can be more informative than scrutinizing one CpG at a time, as it may, for example, help in identifying co-methylation patterns and regions that may be important from a population-epigenetic perspective^66^. Accordingly, we examined regions of 50 kb around each significant CpG detected in our XWAS. This led to the identification of clusters of positively correlated CpGs often mapping to a promoter region (see Figure 4). We also observed clusters of CpGs within gene body regions, such as cg13866977 which was positively correlated with 16 other CpGs across the *AMOT* region (Figure 5). Overall, the patterns of co-methylation in promoter and gene-body regions were anti-correlated between one another, which is as expected and consistent with gene expression patterns typical for these regions^67;68^. To illustrate, three CpGs within a promoter region near *EIF2S3* were highly positively correlated with cg25034591 (Figure 4), in addition to a cluster of positively correlated CpGs within the *EIF2S3* gene body. However, this cluster was *negatively* correlated with cg25034591. The co-methylation analysis of the significant findings in the boys-only XWAS (Figure 6) indicated that both significant CpGs are located within a cluster of highly correlated CpGs and are also part of a DMR located at chr X:84,189,179-84,189,658 (GRCh37) that harbours four CpGs.

**Figure 4:**
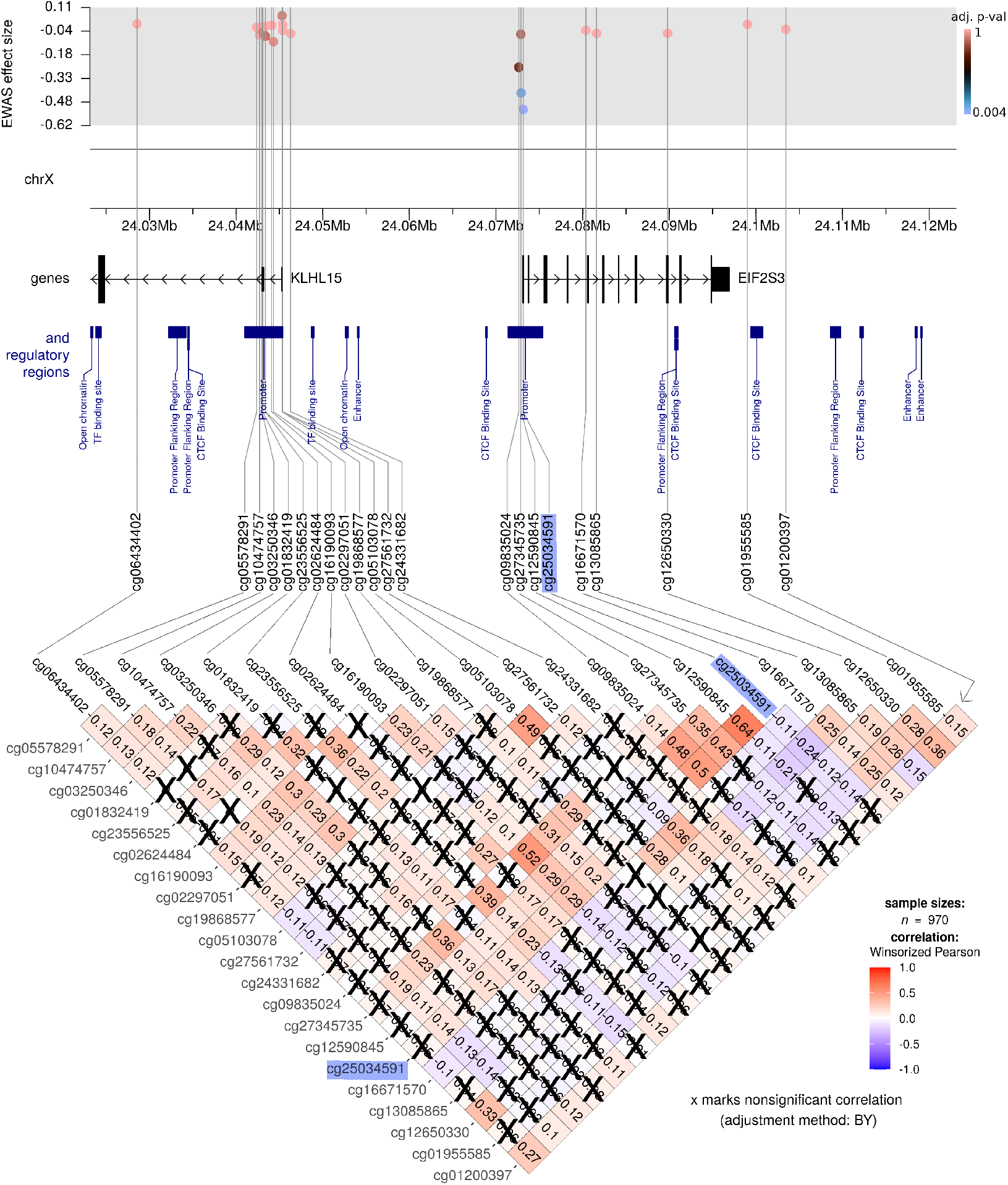
Co-methylation patterns in the region flanking the most significant XWAS finding (cg25034591; highlighted in blue) in the girls-only analysis. The top part of the figure shows the effect sizes and FDR-adjusted p-values for the CpGs within 50 kb of cg25034591. The X chromosome coordinates are provided directly underneath. The middle part of the plot shows the location of genes and regulatory regions based on GRCh37 annotations. The bottom part of the figure shows the DNAm correlation matrix, where the color gradient indicates the strength and direction of correlation of DNAm level for each pair of CpGs. Note that the correlation coefficients are provided inside each matrix element (each diamond), and nonsignificant correlations are crossed out.

**Figure 5:**
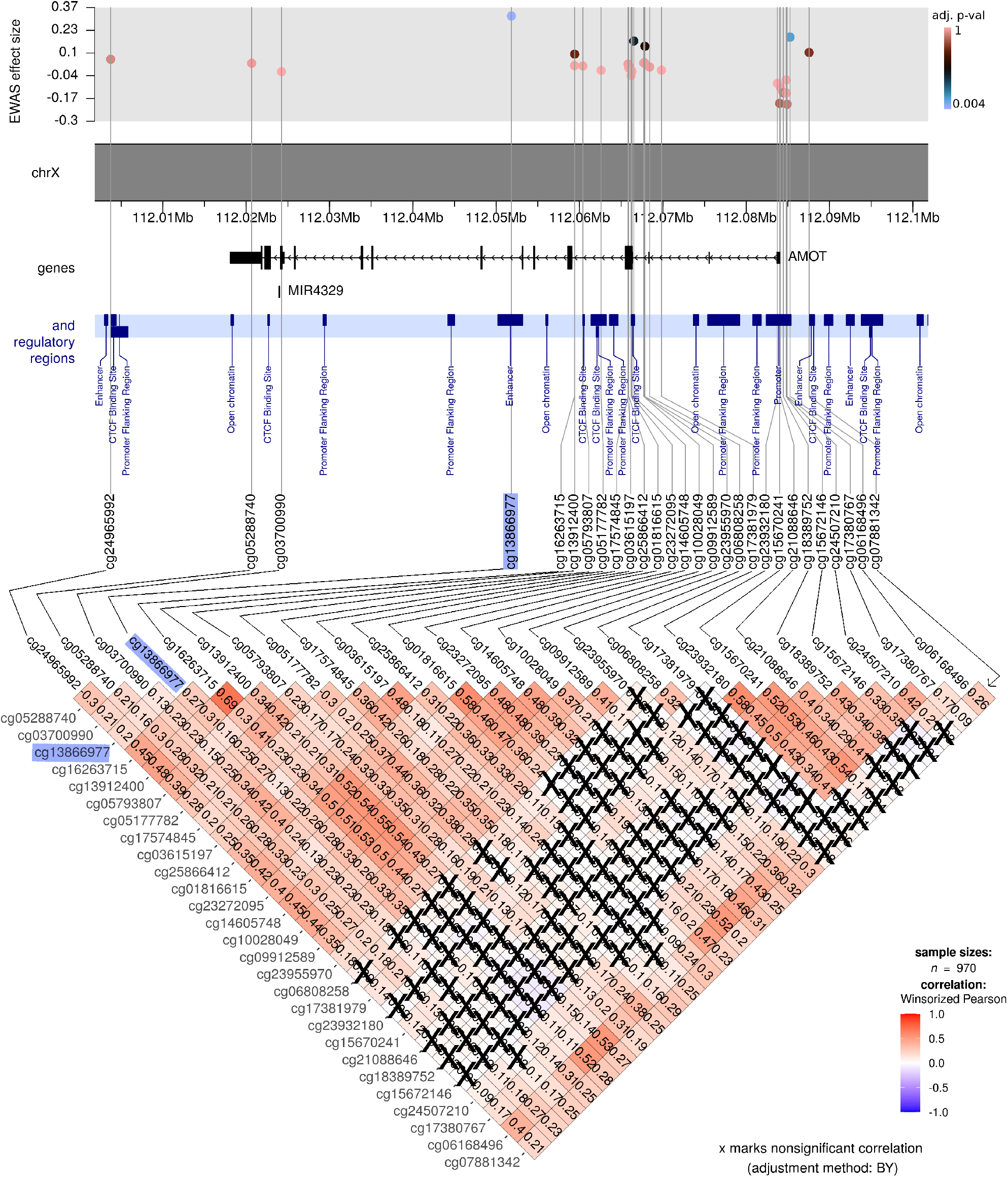
Co-methylation patterns among CpGs within 50 kb of cg13866977 (highlighted in blue). The rest of the figure legend is similar to that of Figure 4 above and will therefore not be repeated here or in the remaining co-methylation figures below.

**Figure 6:**
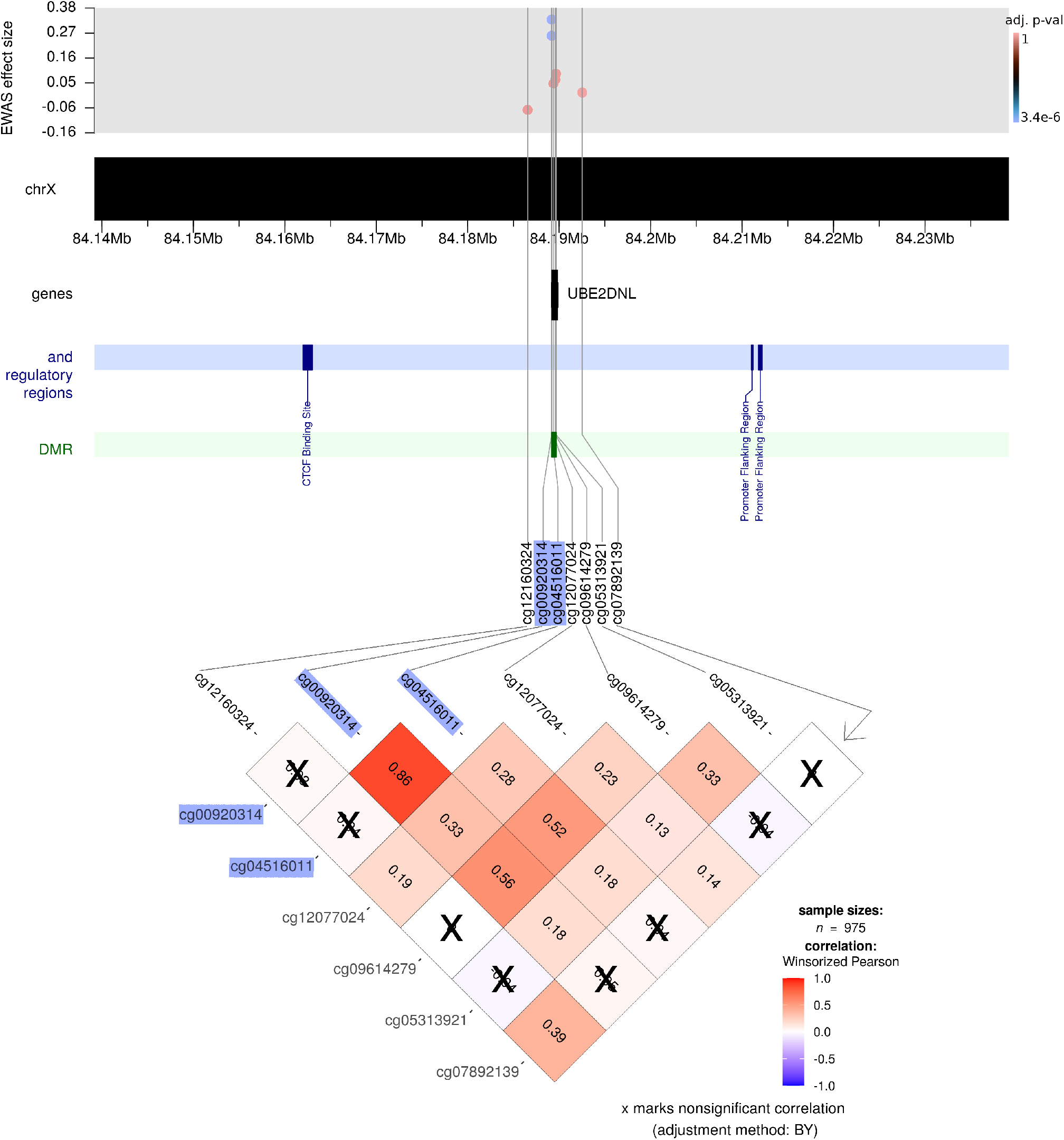
Co-methylation patterns in the area flanking the most significant XWAS finding in the boys-only analysis (cg00920314; highlighted in blue). As with the co-methylation figures for the girls-only analyses, the top part of the plot shows the effect sizes and p-values for the CpGs within 50 kb of cg00920314, which also includes the next most significant CpG, cg04516011 (also highlighted in blue). In addition, the dark-green bar in the plot indicates a DMR in this region.

### 2.4 DMR analysis in ART and non-ART newborns of the MoBa cohort

We identified 12 significant DMRs in the girls-only analysis and three in the boys-only analysis (main model, Figure 7). We considered a DMR as being statistically significant if it contained three or more CpGs and had an FDR-adjusted p-value < 0.01. The number of significant DMRs varied only slightly between the main and the adjusted model (see Supplementary Figures S4–S6); notably, we found eight DMRs in the girls-only analysis and two in the boys-only analysis that were shared across all the models tested. The majority of these DMRs were located in promoter regions. See the Supplementary Document 2 for more details as well as the Github repository for all the results. In one instance, a DMR included a CpG that was significantly associated with the ART phenotype in boys (Figure 6). Overall, however, there was only one instance where the DMRs in boys and girls were near each other (Figure 8, panel A).

**Figure 7:**
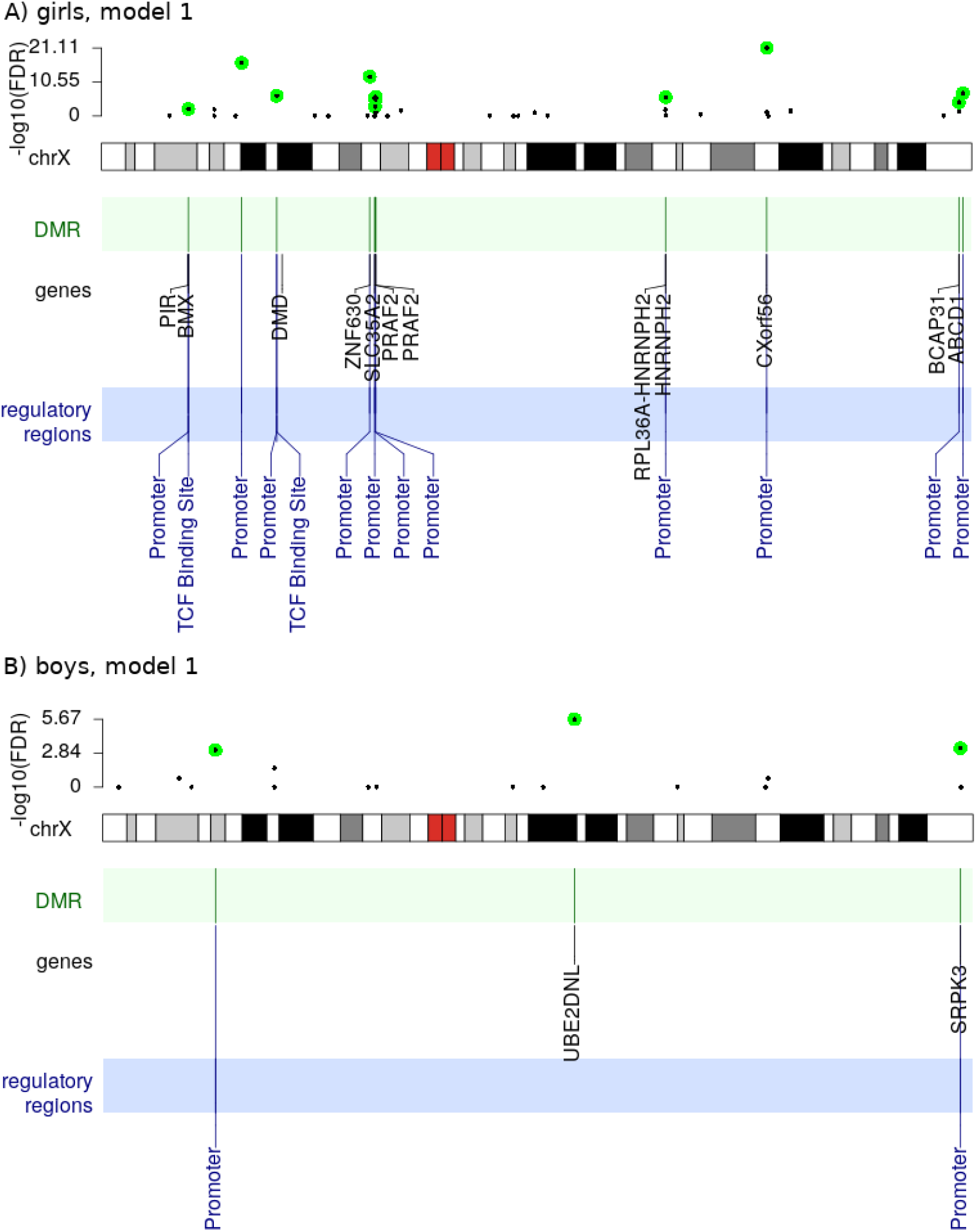
Location of DMRs on the X chromosome. The girls-only analysis is shown in panel A and the boys-only analysis in panel B. Note that we only show the results of the main model (**Model 1**). The top part of each panel shows p-values for all the DMRs that contain at least three CpGs. The FDR-adjusted p-values < 0.01 are marked in green. The bottom part of each panel marks genes and regulatory regions harbored by the significant DMRs.

**Figure 8:**
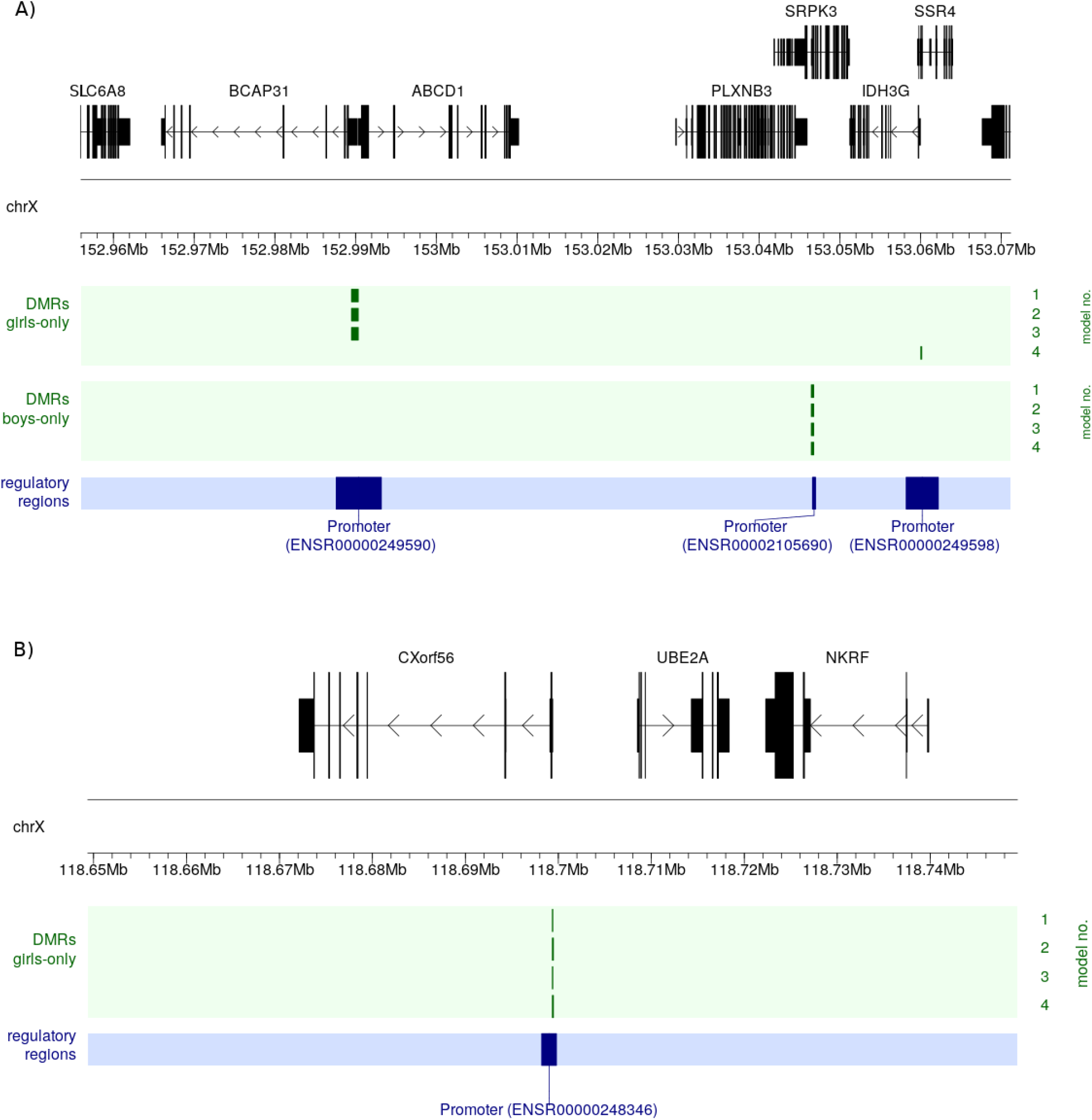
Zooming in on genomic features around selected significant DMRs. A) These significant DMRs detected in the girls-only and boys-only analyses where the only ones localized in the vicinity of each other; the DMRs’ positions are marked by green vertical bars in four lines, where each line corresponds to a specific statistical model (**Models 1-4**; right-hand side). B) The most significant DMR detected in the girls-only analyses.

### 2.5 Testing for replication in an independent cohort

For replication purposes, we ran an analogous model to our main model on an independent dataset from the Australian CHART study (https://lifecourse.melbournechildrens.com/cohorts/art/, see Figure 1 and Methods). Despite the substantially smaller sample size of the CHART cohort (149 ART and 58 non-ART newborns), the results did point to possible associations between ART status and DNAm at CpGs in *EIF2S3* and *AMOT* for girls (Figure S7). Moreover, there was a 401 bp-long DMR in *UBE2DNL* in the boys-only analyses, which contained four hypomethylated probes. Finally, we observed significant differences in DNAm levels at cg04516011 and cg00920314 in boys (Figure S8), which were also identified in the larger discovery (MoBa) sample.

## 3 Discussion

We investigated differences in DNAm levels on the X chromosome between newborns conceived through ART and those conceived naturally. Equipped with the largest collection of ART trios to date, we searched for DNAm differences at single-CpG sites as well as in regions, and ran four separate models to check for the effect of various potential confounders, including parental DNAm. Additionally, we replicated the analysis using the main model on a smaller, but independent, cohort of ART newborns (CHART).

### 3.1 Characteristics of the ART and non-ART participants in the MoBa cohort

The ART parents in our study were older than the non-ART parents, and the ART newborns weighed less than the non-ART newborns. Both of these observations are consistent with previous findings^69–73^. Furthermore, fewer of the ART mothers smoked during pregnancy than the non-ART mothers, but, intriguingly, a higher proportion of the ART mothers were past smokers. The observation that more ART mothers were past smokers is noteworthy in light of previous findings of a link between smoking and impaired fertility in both men and women^74^. Furthermore, in a meta-analysis of 21 studies^75^ significant associations were found between smoking at the time of ART treatment and lower success rate for a number of clinical outcomes of ART. Specifically, smoking was associated with lower odds of live birth per cycle, lower odds of clinical pregnancy per cycle, higher odds of spontaneous miscarriage, and higher odds of ectopic pregnancy.

### 3.2 Significant sex-specific DNAm differences in ART and non-ART newborns

The results of our current XWASs of the MoBa data showed significant sex-specific DNAm differences in ART and non-ART newborns. These differences remained significant even after adjusting for several confounders known to be associated with cord-blood DNAm. The results also revealed more differentially methylated CpGs and DMRs in girls than boys, with a slightly lower overall X-chromosome-wide methylation in girls and the opposite pattern in boys. This sex-dependent pattern is consistent with a previous XWAS of age-associated DNAm patterns in males and females^59^. The differentially methylated CpGs in our study were mostly located in promoters controlling genes involved in several key developmental processes (e.g., neurodevelopment) and disorders (e.g., intellectual disability and autism).

### 3.3 Differential DNAm at cg25034591 suggests upregulation of several genes involved in transcription and translation processes

In the MoBa cohort, the most significant CpG associated with ART in the girls-only analyses, cg25034591, is located in *EIF2S3* and a promoter region (ensembl ID: ENSR00000245352). This promoter region regulates ten genes (https://www.genecards.org/Search/Keyword?queryString=ENSR00000245352, see Suppl. Table S1), five of which encode a highly interconnected group of proteins with important functions in the regulation of transcription and translation (https://version-11-5.string-db.org/cgi/network?networkId=b4AYVO1yPaIh). The DNAm patterns around cg25034591 form two distinct clusters, one containing a set of positively-correlated downstream CpGs and the other a set of negatively-correlated upstream CpGs (Figure 4). This pattern indicates that cg25034591 does not act alone, but operates in concert with other neighboring CpGs. This result was also supported by the independent data from the CHART study, where two other CpGs in *EIF2S3* displayed marked differences in DNAm in ART versus non-ART girls (Figure S7). These CpGs were not present in the MoBa sample analyses because they had been excluded after quality control.

Mutations in *EIF2S3* cause MEHMO, a rare X-linked syndrome characterized by intellectual disability, epilepsy, hypogonadism, hypogenitalism, microcephaly, and obesity^76–78^. Interestingly, both *EIF2S3* and cg25034591 have been reported to escape XCI^33;59^. We also find evidence for this in our data; notably, the *β*-values for DNAm at cg25034591 were within the range 0.00009-0.018 in ART-conceived girls and within 0.00016-0.032 in those naturally-conceived. It is thus plausible that ART interferes with the escape of XCI at this CpG, leading to an upregulation of genes controlled by the promoter ENSR00000245352.

### 3.4 Interpreting the relevance of the findings in the context of ART

The second most significant CpG in girls, cg13866977, lies within a regulatory region and an intron of *AMOT*. This CpG was originally annotated to a region defined as ‘enhancer’ in the GRCh37 (hg19) version of the genome, but was subsequently changed to ‘promoter flanking region’ in the newer GRCh38 (hg38) genome build (ensembl regulatory ID: ENSR00000912938). It is not unusual for the definition and location of an annotation to change from one genome version to another, especially when the distinction between a promoter and an enhancer may become blurred as a result of sharing several properties and functions^79^. A perhaps more suitable annotation for cg13866977 in this case would have been ‘transcription regulatory element’. Furthermore, GeneHancer^80^ lists this regulatory region as a putative enhancer (https://www.genecards.org/Search/Keyword?queryString=ENSR00000912938) for four genes, one of which is *AMOT* (Suppl. Table S1). *AMOT* is a member of the motin family of angiostatin-binding proteins. This gene is especially relevant for ART since it is expressed in placental vessels and the endothelial cells of capillaries, with reported links to premature births^81^. Nevertheless, interpreting the relevance of this finding in the context of ART is not straightforward.

The above-mentioned promoter, ENSR00000912938, is particularly active in six different types of tissues, including the placenta. However, there is no evidence of its activity in cord blood. These observations are based on the ensembl visualization of experimental data showing various histone marker states and DNase1 activity for this promoter (http://www.ensembl.org/Homo_sapiens/Regulation/Summary?db=core;fdb=funcgen;r=X:112806973-112809972;rf=ENSR00000912938). Furthermore, according to the Genotype-Tissue Expression (GTEx) database^82^, neither *AMOT* nor ‘LHFPL tetraspan subfamily member 1’ (*LHFPL1*; another protein-coding gene controlled by this regulatory region) is transcribed in blood, which is paradoxical given that the DNAm data in both the MoBa and the CHART cohort were generated from newborn’s cord blood.

Our results also showed that the DNAm level at cg13866977 was close to 1.0 in boys, implying that the cytosine at this site is fully methylated. In girls it was mostly above 0.7 (Figure 9). Since DNAm signals mainly reflect the level of transcription, we investigated whether transcription factors (TFs) predicted to bind to cg13866977 preferentially bind to the unmethylated or methylated sequence. The output of the search in JASPAR and MeDReaders indicated that none of the seven TFs bind to the methylated sequence (Supplementary Data 3, also available online in the Github repository). This suggests that high methylation at this CpG might signal the inactivation of this regulatory region. Moreover, the methylation state was higher among girls conceived by ART than those conceived naturally (effect size = 0.32). The effect size did not change appreciably when we adjusted for parental DNAm at this site (effect size = 0.33). Again, the independent dataset from the CHART study showed a similar trend of association (Figure S7), except for two other CpGs, cg05177782 and cg09912589, that are positively correlated with cg13866977 (see Figure 5). Although these results suggest that the regulatory region within *AMOT* is less active after the ART procedure in girls, the specific function of this activation remains to be elucidated.

**Figure 9:**
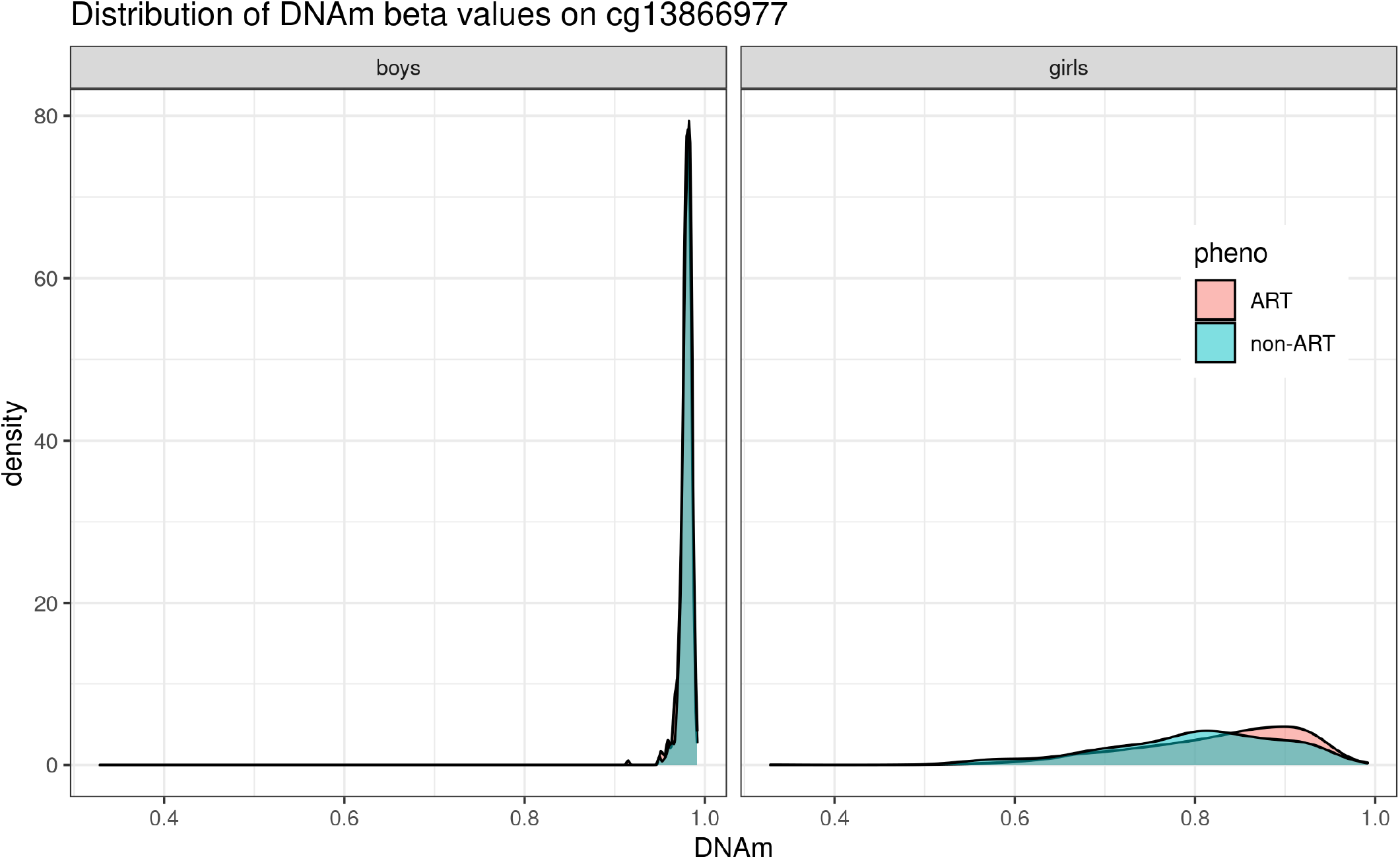
The distribution of methylation β values at cg13866977 differs between girls and boys. This CpG is located within a regulatory region and the *AMOT* gene. There is a significant difference in methylation level between girls conceived by ART and girls conceived naturally.

### 3.5 Differential DNAm in boys point to a pseudogene with unknown function

The most significant CpGs in the boys-only analysis, cg00920314 and cg04516011, were both located within *UBE2DNL*. The NCBI gene database (https://www.ncbi.nlm.nih.gov/gene) classifies *UBE2DNL* as a pseudogene, but the ExpressionAtlas database^83^ (https://www.ebi.ac.uk/gxa/) reports that it is expressed in testes. According to UniProt, *UBE2DNL* is inactive because it lacks a cytosine in the active site (https://www.uniprot.org/uniprotkb/Q8IWF7). There is mounting evidence pointing to pseudogenes as playing important roles in gene regulation instead of just being evolutionary relics of inactive genes^84;85^. This notion has garnered additional support through the application of high-throughput sequencing technologies enabling genome-wide characterizations of pseudogenes^86–88^. Similar to our findings, several of the transcribed pseudogenes identified in an previous study by Zheng *et al*.^86^ were also found to be either transcribed exclusively in testes or were particularly active in those tissues. This pattern of testis-specific pseudogene transcription has also been reported by others^89;90^. The association with *UBE2DNL* in our data also appears to be credible for three reasons: *1)* the CpGs remain significant even after adjusting for covariates (see Supplementary Document 1), *2)* there is a highly significant DMR in this region, and *3)* the data from the independent CHART cohort showed a DMR in this pseudogene (data not shown) and differential DNAm at these two CpGs in ART vs. non-ART newborns (Figure S8).

### 3.6 DMRs co-located with genes involved in key developmental processes

The most significant DMR (chrX:118,699,347–118,699,412 in GRCh37) in the girls-only analyses of the MoBa cohort is located within the promoter of three genes (see Figure 8B and Suppl. Table S2 for details). This promoter is active in cord blood, and the genes linked to this promoter are important for immune response, mitochondrial processes, and chromosome segregation (see the references in the Suppl. Table S2). Specifically, one of the genes controlled by this promoter is ‘STING1 ER exit protein 1’ (*STEEP1*, previously called CXorf56, Figure 8B). Mutations in *STEEP1* cause X-linked intellectual disability and other neurological disorders^91;92^. The DMR encompassing *STEEP1* was also found to have a lower level of DNAm in ART compared to non-ART newborns, suggesting that the expression of these genes might be up-regulated by ART. The link with *STEEP1* needs to be verified in other similar cohorts.

Another significant DMR (chrX:152,989,492–152,990,345 in GRCh37) in the girls-only analyses was co-located with a promoter (ensembl ID: ENSR00000249590, see Figure 8A), which, according to a search in GeneHancer DB, is either a putative promoter or enhancer for nine genes (see Suppl. Table S2 for more details). Six of these genes encode proteins that form part of a network, according to the results of text mining and co-expression arrays (STRING DB, https://version-11-5.string-db.org/cgi/network?networkId=bWgRg6ih0nDV). Deletions or duplications in many of these genes have been reported to cause different impairments and diseases^93–99^, with autism featuring prominently among these clinical manifestations. Right next to this DMR (chrX:152,989,492-152,990,345 in GRCh37), the boys-only analysis revealed another significant DMR (chrX:153,046,451-153,046,767 in GRCh37) co-located with a promoter (ensembl ID: ENSR00002105690, see Fig. 8A) that is active in cord blood. This promoter regulates four genes that are important in the developmental processes of various tissues, including neurons (Suppl. Table S2). These indirect connections with autism and neurodevelopment are particularly noteworthy, given previous reports indicating that neurodevelopmental outcomes differ in children conceived by ART^100;101^, but not always^102;103^.

Lastly, we also checked for any common features among all the genes that co-localized with all the significant DMRs. The STRING protein-protein interaction database (https://version-11-5.string-db.org/cgi/network?networkId=b1fEsljWdfBy) indicates that five of the 13 genes found in all the significant DMRs in the girls-only analysis are involved in X-linked monogenic diseases (DOID:0050735, https://diseases.jensenlab.org/).

### 3.7 Strengths and weaknesses

A major strength of our study is the large size of the MoBa sample, enabling a more powerful exploration of questions related to ART and infertility. Additionally, the trio design enabled adjusting for parental DNAm in the regression models, which is essential to rule out other underlying parental characteristics, e.g., parental infertility, as a possible reason for the observed associations in the newborns. Another strength is the mandatory reporting of any use of ART to the Norwegian Medical Birth Registry, including the specifics of the ART procedure used to achieve pregnancy. This ensures virtually complete case ascertainment. Combined with the detailed data from questionnaires on relevant covariates, the depth of information on these trios is unparalleled. Furthermore, the DNAm data were generated on the more comprehensive Illumina EPIC array, which is a significant technical leap over its predecessors (Illumina’s 27K and 450K Beadchips) in terms of its genomic coverage of regulatory elements, reliability, and reproducibility^104^. One shortcoming of our study is the lack of a well-powered replication cohort with which to compare and validate our findings. EWASs of ART have been far and few between. To our knowledge, the only available dataset was CHART – a small cohort from Australia. Nonetheless, the results of the main XWAS model in the CHART cohort showed the same trends as observed in the MoBa cohort, despite CHART being significantly smaller and stemming from a different population than MoBa.

## 4 Conclusions

To summarize, our results showed that, for newborns conceived with the help of ART, there were more differentially methylated CpGs and DMRs in girls than boys, with a slightly lower genome-wide methylation in girls and the opposite pattern in boys. Adjustment for several confounders known to be associated with cord-blood DNAm did not affect the associations, nor did adjustment for parental DNAm, which makes it less likely that parental characteristics (including parental infertility) were responsible for the observed associations in the newborns. Moreover, our downstream bioinformatic analyses revealed that several of the identified genes were expressed in tissues that are relevant for ART and sex. A number of the genes are associated with neurodevelopment and intellectual disability, which is consistent with previous reports of significant differences in neurodevelopment between newborns conceived by ART and those conceived naturally. More generally, our study fills an important knowledge gap in that it provides an easily adaptable analytic pipeline to investigate the contributions of X-linked CpGs to subfertility and other traits. Its application to the reanalysis of previously published EWASs, such as those in the EWAS Open Platform^105^ and the GEO repository, may facilitate the discovery of additional genes and loci that might have been missed by focusing solely on autosomal CpGs.

## 5 Methods

### 5.1 Discovery cohort – MoBa

MoBa is a large population-based pregnancy cohort study in which pregnant women were recruited across Norway from 1999 through 2008^63^. Fathers were invited from 2001 onward. The participation rate was 41% among the MoBa mothers. Overall, MoBa includes 114 000 children, 95 000 mothers and 75 000 fathers. Blood samples were initially drawn from the parents at approximately 18 weeks of gestation, and later from the mother and the umbilical cord after delivery^106^. The current analyses were done on a subset of MoBa data, generated in the ‘Study of Assisted Reproductive Technology’ (START) project^62^. The current MoBa dataset included 963 trios in which the newborn was conceived using ART and 982 randomly sampled trios in which the newborn was conceived naturally (i.e., by coitus). DNAm in both of these groups of trios was measured using the Illumina Infinium Methylation EPIC BeadChip (Illumina, San Diego, USA) which houses ~850 000 CpG sites. The inclusion criteria consisted of all of the following: *1)* the child was born in the period 2001-2009, *2)* the child was a singleton newborn with a record in the Norwegian Medical Birth Registry, *3)* the mother filled out and returned the first MoBa questionnaire at around week 17 of gestation, and *4)* blood samples were available for the whole trio (child, mother, and father).

In Norway, fertility clinics are mandated to report any ART conception to the national birth registry. We defined ART as “any ART” (excluding intrauterine insemination) and coded it as a binary variable (ART vs. non-ART). As information on the ART procedure was missing for 79 of the trios, these were excluded from the analysis.

### 5.2 DNAm measurements in the discovery cohort

DNA samples from the ART and non-ART trios in the MoBa cohort were shipped to the Institute of Life & Brain Sciences at the University of Bonn, Germany, for further sample processing and measurement of DNAm on the Illumina Infinium Methylation EPIC Bead-Chip platform (Illumina, San Diego, USA). Extensive details regarding the quality control (QC) pipeline used for data cleaning have been provided in our previous work^62^. Briefly, we established a QC pipeline based on the `RnBeads` package^107^ using the statistical programming language R^108^. Cross-hybridizing probes and probes in which the last three bases overlapped with a SNP were removed from the analyses. Additionally, probes with a detection p-value above 0.01 were used in the greedycut algorithm to remove unreliable probes and samples. This procedure minimizes the false positive rate and maximizes the sensitivity when the retained measurements are considered as prediction for the reliable ones. The remaining DNAm data were corrected for background noise using the `enmix.oob` function^109^.

We extracted DNAm data on the X chromosome only and applied BMIQ^110^ to normalize the Type I and Type II probes^111^. We then checked for multimodality of DNAm per CpG for girls and boys separately using the `gaphunter` function in the `minfi` R package^112;113^. Crucially, the QC functions applied to the data did not combine any information across samples, which is essential to keep the analyses separate for males and females due to their distinct modalities. The total number of probes on the X chromosome remaining for the current analyses was 16,841, out of the initial 19,090 X chromosome probes present on the EPIC array.

Figure 1 provides an overview of the analytic pipeline and study population, and the sections below provide additional details.

### 5.3 Statistical analyses in the discovery cohort

#### Regression models

In preparation for the XWAS of the MoBa cohort, we used the `logit2()` function from R package `minfi`^113^ to transform *β*-values for DNAm into M-values, since M-values are more amenable to statistical tests^114^. Four regression models were fit for boys and girls separately to estimate the difference in methylation levels between the ART and non-ART newborns. This stratification by sex is necessary because of the distinctly different overall DNAm profiles for girls and boys on the X chromosome. In previously published studies, a number of variables were reported to be associated with DNAm in cord blood and with the use of ART, which includes mother’s age, smoking status, BMI, and whether she was primiparous. These variables were included in the model as potential confounders, i.e., `CpG` ~ `ART + maternal age + maternal smoking + maternal BMI + primiparity` (referred to as the ‘main model’; see also Figure 1). Although all samples were randomly placed on the bisulfite conversion plates before measuring DNAm, the regression model also included plate ID as a random effect to adjust for batch effects. As DNAm levels associated with parental infertility may confound the XWAS results in the newborns, we ran additional models where we adjusted for maternal methylation in the boys-only analysis and for both maternal and paternal methylation in the girls-only analysis (referred to as the ‘adjusted model’). Moreover, we extended each of the two aforementioned models by including further adjustments for gestational age and birthweight (see Figure 1). Linear mixed models were implemented using the *rint*.*reg* function in the R package *Rfast* ^115;116^.

#### Controlling for inflation of the test statistics

Supplementary Figure S1 depicts the density curves of the DNAm values (*β*-values) in the newborns according to sex and type of probe on the Illumina EPIC array. The methylation patterns are distinctively different in males and females, as has also been reported by other studies (e.g.,^59^). The middle portion of the distribution for females typically exhibits a bump, as a consequence of XCI, whereas males exhibit higher densities at the opposite ends of the distribution. As females have two copies of the X chromosome, and one copy is silenced through XCI, the distribution of the average DNAm is flatter in females than males.

As pointed out by several reports^117–119^, large-scale hypothesis testing of high-dimensional data (e.g., those stemming from a GWAS, EWAS, or XWAS) may be prone to heavily inflated type I error when using the theoretical null distribution to assess the significance of the p-values. We, therefore, used the R package BACON^119^ to re-scale the raw z-statistics from the XWAS. BACON is a Bayesian method that controls the false positive rate and accounts for potentially poorly calibrated test statistics while preserving statistical power. We chose BACON over competing methods because it is flexible and can handle a larger proportion of true associations^119;120^. As can be seen in Supplementary Figure S2, BACON reduced inflation substantially in girls but had a negligible effect in boys. After this correction, we applied a false discovery rate (FDR) < 0.01 to select CpGs that were significantly associated with ART in our sample.

### 5.4 Consistency of significant findings in the discovery cohort

We applied a bootstrapping scheme to the XWAS results to evaluate the consistency with which a significant CpG was identified as being significant. We created 1000 bootstrap samples with replacement separately for girls and boys, ensuring an equal proportion of ART and non-ART cases as in the original MoBa dataset. We then reran the analysis using the same main model for each of the bootstrap samples and determined the proportion of times each CpG was found to be significant (using the same significance threshold as previously).

### 5.5 Co-methylated CpGs and DMR detection in the discovery cohort

We retrieved the annotation tracks from Ensembl BioMart^121^ (http://www.ensembl.org) using the R package `biomaRt`^122;123^ and generated a regional plot of the association results. This regional plot was subsequently combined with a co-methylation (correlation) plot of neighboring CpG sites flanking the significant CpGs. The correlation of DNAm values was calculated and plotted using `ggstatsplot`^124^. The rationale is that, if the biological functions of two CpGs are correlated, their DNAm levels are expected to change in the same way between ART and non-ART samples.

We chose the `dmrff` R package^125^ to identify differentially methylated regions (DMRs). This choice was based on the results of a recent paper demonstrating the superior performance of `dmrff` to four frequently used methods for DMR detection: DMRcate, comb-p, seqlm, and GlobalP^126^. Finally, the R package `karyoploteR` (part of Bioconductor^127^) was used to visualize genomic features superimposed on a linear representation of the X chromosome^128^.

### 5.6 Downstream bioinformatic analyses in the discovery cohort

The most significant CpGs and DMRs (both at FDR < 0.01) from the above analyses were subjected to a series of downstream bioinformatic analyses to unravel the biological processes that might be influenced by DNAm at these CpGs. Briefly, we searched the MeDReaders database^129^ (http://medreader.org/) and adapted data from Yin *et al*.^130^ to retrieve information about transcription factors (TFs) that preferentially bind to methylated DNA (the table is available in the Github repository). We also used JASPAR2022^131^ tracks in the ensembl genome browser (http://grch37.ensembl.org/Homo_sapiens/) to check which TFs bind to the significant CpGs detected in our XWASs. GeneHancer^80^ was used to find possible targets of promoter and enhancer regions that are co-located with our results. Further, to obtain information on mRNA transcription and protein expression, we searched ExpressionAtlas^132^ (https://www.ebi.ac.uk/gxa/home) and HumanProteinAtlas^133^ (https://www.proteinatlas.org/). Finally, information about gene and protein functions and interactions were gathered via GeneCards^134^, UniProt^135^, and STRING db^136^.

### 5.7 Replication cohort and analysis

For replication purposes, we analyzed the DNAm data in the Australian CHART cohort. CHART consists of 547 adults conceived with the use of IVF and 549 naturally-conceived controls^64;65;137^. In a subsample of 149 ART-conceived and 58 non-ART neonates (see Figure 1), DNAm was measured in DNA isolated from neonatal blood spots (Guthrie spots) using the Illumina EPIC array. Data was pre-processed using the `MissMethyl` R package^138^ and low quality and cross-reactive probes were removed from analysis^15^. Cell composition was estimated using the Bakulski cord blood cell reference method^139^. Maternal smoking during pregnancy was predicted using a DNA Methylation score^140^. Linear regression modelling was performed using the `limma` R package^141^, with the model: `CpG` ~ `ART + maternal smoking + sentrix ID + sample well + sentrix position + sample plate`. The analyses were run separately for boys and girls.

## 6 Declarations

### 6.1 Ethics approval and consent to participate

The establishment of MoBa and the initial data collection were based on a license from the Norwegian Data Protection Agency and additional approvals from the Regional Committees for Medical and Health Research Ethics (REK) in Norway. The MoBa cohort is now based on regulations in the Norwegian Health Registry Act. The current study was approved by the Southeast branch of REK in Norway (reference number 2017/1362).

The Australian CHART study was approved by the Human Research Ethics Committee (RCH HREC Project 33163) of the Royal Children’s Hospital, Melbourne, Australia.

### 6.2 Availability of data and materials

The MoBa and START data are available from the NIPH, but restrictions apply regarding the availability of these data, which were originally used under specific approvals for the current study and are therefore not publicly available. Data are, however, available from the authors upon reasonable request and after approval by relevant authorities and the NIPH.

The CHART data have been deposited in the Gene Expression Omnibus repository, under accession number GSE131433.

The scripts for all the above analyses as well as all the results of calculations are available in the GitHub repository at https://github.com/folkehelseinstituttet/X-factor-ART.

### 6.5 Authors’ contributions

HEN and CMP pre-processed raw DNAm data and performed quality control.

JR and HEN ran analyses and interpreted the results, as well as wrote the manuscript and prepared the code and data for sharing in the Github repository.

AJ formulated the research question, drafted the manuscript, and was a major contributor to the manuscript overall.

JB, WRPD, YL, MCM, KLH, MG, HKG, RL, PM, SEH helped shape the final version of the manuscript.

JB, AJ, JR, and HEN were heavily involved in discussing the results.

MG, CMP, WRPD, and HKG checked the validity of the statistical methods used.

BN and RS performed replication analyses and created figures, as well as contributed to the relevant parts of the manuscript.

PM and SEH secured funding necessary for obtaining raw DNA methylation data. All authors read and approved the final manuscript.

## 6.6 Acknowledgements

MoBa is supported by the Norwegian Ministry of Health and Care Services and the Ministry of Education and Research. We are grateful to all the participating families who take part in this ongoing cohort study.

