## Supplementary documents 1 and 2 for "The X-factor in ART: does the use of Assisted Reproductive Technologies influence DNA methylation on the X chromosome?": Supplementary_document1.html

| **Most significant findings (FDR < 0.01)** | | | | | | | | | | |
| --- | --- | --- | --- | --- | --- | --- | --- | --- | --- | --- |
| *(grouped by sex and model)* | | | | | | | | | | |
| **CpG info** | | **EWAS results** | | **Regulatory regions**^1^ | | | **Genes**^1^ | | | |
| --- | --- | --- | --- | --- | --- | --- | --- | --- | --- | --- |
| CpG ID | Position (chrX)^2^ | Effect size^3^ | FDR | Ensembl ID | Region type | Position^2^ | Ensembl ID | Description | Gene ID | Position^2^ |
| Girls - Model 1 | | | | | | | | | | |
| cg25034591 | 24,073,134 | −0.52 | 3.77 × 10^−3^ | ENSR00000245352 | Promoter | chrX:24,071,400-24,075,401 | ENSG00000130741 | eukaryotic translation initiation factor 2, subunit 3 gamma, 52kDa [Source:HGNC Symbol;Acc:3267] | EIF2S3 | chrX:24,072,833-24,096,088 |
| cg13866977 | 112,051,826 | 0.32 | 3.77 × 10^−3^ | ENSR00001768065 | Enhancer | chrX:112,050,201-112,053,200 | ENSG00000126016 | angiomotin [Source:HGNC Symbol;Acc:17810] | AMOT | chrX:112,017,731-112,084,043 |
| cg26175661 | 114,826,443 | 0.28 | 5.71 × 10^−3^ | NA | NA | NA | ENSG00000102024 | plastin 3 [Source:HGNC Symbol;Acc:9091] | PLS3 | chrX:114,795,501-114,885,181 |
| Girls - Model 2 | | | | | | | | | | |
| cg25034591 | 24,073,134 | −0.54 | 1.48 × 10^−3^ | ENSR00000245352 | Promoter | chrX:24,071,400-24,075,401 | ENSG00000130741 | eukaryotic translation initiation factor 2, subunit 3 gamma, 52kDa [Source:HGNC Symbol;Acc:3267] | EIF2S3 | chrX:24,072,833-24,096,088 |
| cg13866977 | 112,051,826 | 0.33 | 1.48 × 10^−3^ | ENSR00001768065 | Enhancer | chrX:112,050,201-112,053,200 | ENSG00000126016 | angiomotin [Source:HGNC Symbol;Acc:17810] | AMOT | chrX:112,017,731-112,084,043 |
| Girls - Model 3 | | | | | | | | | | |
| cg25034591 | 24,073,134 | −0.52 | 4.46 × 10^−3^ | ENSR00000245352 | Promoter | chrX:24,071,400-24,075,401 | ENSG00000130741 | eukaryotic translation initiation factor 2, subunit 3 gamma, 52kDa [Source:HGNC Symbol;Acc:3267] | EIF2S3 | chrX:24,072,833-24,096,088 |
| cg13866977 | 112,051,826 | 0.32 | 4.46 × 10^−3^ | ENSR00001768065 | Enhancer | chrX:112,050,201-112,053,200 | ENSG00000126016 | angiomotin [Source:HGNC Symbol;Acc:17810] | AMOT | chrX:112,017,731-112,084,043 |
| cg26175661 | 114,826,443 | 0.28 | 7.50 × 10^−3^ | NA | NA | NA | ENSG00000102024 | plastin 3 [Source:HGNC Symbol;Acc:9091] | PLS3 | chrX:114,795,501-114,885,181 |
| Girls - Model 4 | | | | | | | | | | |
| cg25034591 | 24,073,134 | −0.54 | 1.72 × 10^−3^ | ENSR00000245352 | Promoter | chrX:24,071,400-24,075,401 | ENSG00000130741 | eukaryotic translation initiation factor 2, subunit 3 gamma, 52kDa [Source:HGNC Symbol;Acc:3267] | EIF2S3 | chrX:24,072,833-24,096,088 |
| cg13866977 | 112,051,826 | 0.32 | 1.72 × 10^−3^ | ENSR00001768065 | Enhancer | chrX:112,050,201-112,053,200 | ENSG00000126016 | angiomotin [Source:HGNC Symbol;Acc:17810] | AMOT | chrX:112,017,731-112,084,043 |
| Boys - Model 1 | | | | | | | | | | |
| cg00920314 | 84,189,177 | 0.33 | 3.39 × 10^−6^ | NA | NA | NA | ENSG00000229547 | ubiquitin-conjugating enzyme E2D N-terminal like (pseudogene) [Source:HGNC Symbol;Acc:28656] | UBE2DNL | chrX:84,189,157-84,189,896 |
| cg04516011 | 84,189,179 | 0.26 | 4.96 × 10^−6^ | NA | NA | NA | ENSG00000229547 | ubiquitin-conjugating enzyme E2D N-terminal like (pseudogene) [Source:HGNC Symbol;Acc:28656] | UBE2DNL | chrX:84,189,157-84,189,896 |
| Boys - Model 2 | | | | | | | | | | |
| cg00920314 | 84,189,177 | 0.32 | 4.61 × 10^−6^ | NA | NA | NA | ENSG00000229547 | ubiquitin-conjugating enzyme E2D N-terminal like (pseudogene) [Source:HGNC Symbol;Acc:28656] | UBE2DNL | chrX:84,189,157-84,189,896 |
| cg04516011 | 84,189,179 | 0.26 | 4.61 × 10^−6^ | NA | NA | NA | ENSG00000229547 | ubiquitin-conjugating enzyme E2D N-terminal like (pseudogene) [Source:HGNC Symbol;Acc:28656] | UBE2DNL | chrX:84,189,157-84,189,896 |
| Boys - Model 3 | | | | | | | | | | |
| cg00920314 | 84,189,177 | 0.32 | 9.33 × 10^−6^ | NA | NA | NA | ENSG00000229547 | ubiquitin-conjugating enzyme E2D N-terminal like (pseudogene) [Source:HGNC Symbol;Acc:28656] | UBE2DNL | chrX:84,189,157-84,189,896 |
| cg04516011 | 84,189,179 | 0.25 | 9.33 × 10^−6^ | NA | NA | NA | ENSG00000229547 | ubiquitin-conjugating enzyme E2D N-terminal like (pseudogene) [Source:HGNC Symbol;Acc:28656] | UBE2DNL | chrX:84,189,157-84,189,896 |
| Boys - Model 4 | | | | | | | | | | |
| cg00920314 | 84,189,177 | 0.31 | 1.15 × 10^−5^ | NA | NA | NA | ENSG00000229547 | ubiquitin-conjugating enzyme E2D N-terminal like (pseudogene) [Source:HGNC Symbol;Acc:28656] | UBE2DNL | chrX:84,189,157-84,189,896 |
| cg04516011 | 84,189,179 | 0.25 | 1.15 × 10^−5^ | NA | NA | NA | ENSG00000229547 | ubiquitin-conjugating enzyme E2D N-terminal like (pseudogene) [Source:HGNC Symbol;Acc:28656] | UBE2DNL | chrX:84,189,157-84,189,896 |
|  |  |  |  |  |  |  |  |  |  |  |
| --- | --- | --- | --- | --- | --- | --- | --- | --- | --- | --- |
| ^1^ NA = Not applicable | | | | | | | | | | |
| ^2^ genome build GRCh 37 | | | | | | | | | | |
| ^3^ Color shading reflects the magnitude and the sign of the estimated effect - from dark red (strong negative effect), through beige (weak negative effect), and dark blue (strong positive effect). | | | | | | | | | | |
