## Supplementary documents 1 and 2 for "The X-factor in ART: does the use of Assisted Reproductive Technologies influence DNA methylation on the X chromosome?": Supplementary_document2.html

Significant DMR findings, XWAS of ART vs. non-ART


|  |  |  |  |  |  |  |  |  |  |  |  |  |
| --- | --- | --- | --- | --- | --- | --- | --- | --- | --- | --- | --- | --- |
| Model | | DMR info | | | | Regulatory regions | | | Genes | | | |
| group | model | Name | n | estimate | p.adjust | region\_ID | region\_type | region\_position | gene\_ID | gene\_descr | gene\_name | gene\_position |
| Boys | Model 1 | chrX: 20,135,231- 20,135,682 | 6 | 0.64 | 7.98e-04 | ENSR00000245106 | Promoter | chrX:20,134,000-20,135,600 |  |  |  |  |
|  |  | chrX: 84,189,179- 84,189,658 | 4 | 1.04 | 2.13e-06 |  |  |  | ENSG00000229547 | ubiquitin-conjugating enzyme E2D N-terminal like (pseudogene) [Source:HGNC Symbol;Acc:28656] | UBE2DNL | chrX:84,189,157-84,189,896 |
|  |  | chrX:153,046,451-153,046,767 | 5 | 1.04 | 5.66e-04 | ENSR00002105690 | Promoter | chrX:153,046,600-153,047,001 | ENSG00000184343 | SRSF protein kinase 3 [Source:HGNC Symbol;Acc:11402] | SRPK3 | chrX:153,041,867-153,051,187 |
|  | Model 2 | chrX: 20,135,231- 20,135,682 | 6 | 0.64 | 9.57e-04 | ENSR00000245106 | Promoter | chrX:20,134,000-20,135,600 |  |  |  |  |
|  |  | chrX: 84,189,179- 84,189,658 | 4 | 1.05 | 1.32e-06 |  |  |  | ENSG00000229547 | ubiquitin-conjugating enzyme E2D N-terminal like (pseudogene) [Source:HGNC Symbol;Acc:28656] | UBE2DNL | chrX:84,189,157-84,189,896 |
|  |  | chrX:153,046,451-153,046,767 | 5 | 1.00 | 1.96e-03 | ENSR00002105690 | Promoter | chrX:153,046,600-153,047,001 | ENSG00000184343 | SRSF protein kinase 3 [Source:HGNC Symbol;Acc:11402] | SRPK3 | chrX:153,041,867-153,051,187 |
|  | Model 3 | chrX: 13,587,443- 13,587,678 | 5 | 0.42 | 6.79e-04 |  |  |  |  |  |  |  |
|  |  | chrX: 20,135,231- 20,135,682 | 6 | 0.62 | 2.42e-03 | ENSR00000245106 | Promoter | chrX:20,134,000-20,135,600 |  |  |  |  |
|  |  | chrX: 30,671,314- 30,671,410 | 5 | -0.93 | 8.92e-03 | ENSR00000245518 | Promoter | chrX:30,670,200-30,675,201 |  |  |  |  |
|  |  | chrX: 84,189,179- 84,189,658 | 4 | 1.03 | 3.16e-06 |  |  |  | ENSG00000229547 | ubiquitin-conjugating enzyme E2D N-terminal like (pseudogene) [Source:HGNC Symbol;Acc:28656] | UBE2DNL | chrX:84,189,157-84,189,896 |
|  |  | chrX:100,546,291-100,546,519 | 4 | 0.44 | 9.60e-08 | ENSR00001482488 | Promoter | chrX:100,546,001-100,546,600 | ENSG00000102387 | TAF7-like RNA polymerase II, TATA box binding protein (TBP)-associated factor, 50kDa [Source:HGNC Symbol;Acc:11548] | TAF7L | chrX:100,523,241-100,548,059 |
|  |  | chrX:153,046,451-153,046,767 | 5 | 1.03 | 1.07e-03 | ENSR00002105690 | Promoter | chrX:153,046,600-153,047,001 | ENSG00000184343 | SRSF protein kinase 3 [Source:HGNC Symbol;Acc:11402] | SRPK3 | chrX:153,041,867-153,051,187 |
|  | Model 4 | chrX: 13,587,443- 13,587,678 | 5 | 0.44 | 1.18e-04 |  |  |  |  |  |  |  |
|  |  | chrX: 20,135,231- 20,135,682 | 6 | 0.62 | 2.47e-03 | ENSR00000245106 | Promoter | chrX:20,134,000-20,135,600 |  |  |  |  |
|  |  | chrX:153,046,451-153,046,767 | 5 | 0.99 | 3.53e-03 | ENSR00002105690 | Promoter | chrX:153,046,600-153,047,001 | ENSG00000184343 | SRSF protein kinase 3 [Source:HGNC Symbol;Acc:11402] | SRPK3 | chrX:153,041,867-153,051,187 |
| Girls | Model 1 | chrX: 15,511,530- 15,512,019 | 9 | 0.58 | 6.44e-03 | ENSR00000244760 | Promoter | chrX:15,510,000-15,512,201 | ENSG00000087842 | pirin (iron-binding nuclear protein) [Source:HGNC Symbol;Acc:30048] | PIR | chrX:15,402,921-15,511,687 |
|  |  |  |  |  |  |  |  |  | ENSG00000102010 | BMX non-receptor tyrosine kinase [Source:HGNC Symbol;Acc:1079] | BMX | chrX:15,482,369-15,574,652 |
|  |  |  |  |  |  | ENSR00000422673 | CTCF Binding Site | chrX:15,511,801-15,512,200 | ENSG00000087842 | pirin (iron-binding nuclear protein) [Source:HGNC Symbol;Acc:30048] | PIR | chrX:15,402,921-15,511,687 |
|  |  |  |  |  |  |  |  |  | ENSG00000102010 | BMX non-receptor tyrosine kinase [Source:HGNC Symbol;Acc:1079] | BMX | chrX:15,482,369-15,574,652 |
|  |  | chrX: 25,034,241- 25,034,688 | 4 | -0.58 | 3.21e-17 | ENSR00002096583 | Promoter | chrX:25,029,001-25,034,600 |  |  |  |  |
|  |  | chrX: 31,284,529- 31,285,029 | 5 | -0.18 | 5.86e-07 | ENSR00000245550 | Promoter | chrX:31,281,801-31,285,401 | ENSG00000198947 | dystrophin [Source:HGNC Symbol;Acc:2928] | DMD | chrX:31,115,794-33,357,558 |
|  |  |  |  |  |  | ENSR00002097003 | CTCF Binding Site | chrX:31,285,001-31,285,200 |  |  |  |  |
|  |  | chrX: 47,863,437- 47,863,595 | 5 | -0.13 | 6.11e-13 | ENSR00000423313 | Promoter | chrX:47,861,600-47,864,400 | ENSG00000221994 | zinc finger protein 630 [Source:HGNC Symbol;Acc:28855] | ZNF630 | chrX:47,842,756-47,931,025 |
|  |  | chrX: 48,768,940- 48,769,091 | 3 | -0.24 | 2.89e-06 | ENSR00000246526 | Promoter | chrX:48,767,200-48,770,801 | ENSG00000102100 | solute carrier family 35 (UDP-galactose transporter), member A2 [Source:HGNC Symbol;Acc:11022] | SLC35A2 | chrX:48,760,459-48,769,235 |
|  |  | chrX: 48,815,747- 48,815,822 | 3 | -0.13 | 1.39e-03 | ENSR00000423354 | Promoter | chrX:48,812,800-48,817,201 |  |  |  |  |
|  |  | chrX: 48,931,572- 48,931,623 | 3 | -0.11 | 6.43e-06 | ENSR00000423364 | Promoter | chrX:48,929,201-48,932,401 | ENSG00000243279 | PRA1 domain family, member 2 [Source:HGNC Symbol;Acc:28911] | PRAF2 | chrX:48,928,813-48,931,730 |
|  |  |  |  |  |  |  |  |  | ENSG00000250232 | WD repeat domain phosphoinositide-interacting protein 4 [Source:UniProtKB/TrEMBL;Acc:A6NM71] | AF196779.12 | chrX:48,929,385-48,937,546 |
|  |  |  |  |  |  |  |  |  | ENSG00000196998 | WD repeat domain 45 [Source:HGNC Symbol;Acc:28912] | WDR45 | chrX:48,929,385-48,958,108 |
|  |  | chrX: 48,931,699- 48,931,851 | 5 | -0.39 | 1.36e-06 |  |  |  | ENSG00000243279 | PRA1 domain family, member 2 [Source:HGNC Symbol;Acc:28911] | PRAF2 | chrX:48,928,813-48,931,730 |
|  |  |  |  |  |  |  |  |  | ENSG00000250232 | WD repeat domain phosphoinositide-interacting protein 4 [Source:UniProtKB/TrEMBL;Acc:A6NM71] | AF196779.12 | chrX:48,929,385-48,937,546 |
|  |  |  |  |  |  |  |  |  | ENSG00000196998 | WD repeat domain 45 [Source:HGNC Symbol;Acc:28912] | WDR45 | chrX:48,929,385-48,958,108 |
|  |  | chrX:100,663,213-100,663,543 | 3 | -0.50 | 1.50e-06 | ENSR00000423848 | Promoter | chrX:100,660,200-100,666,201 | ENSG00000257529 | RPL36A-HNRNPH2 readthrough [Source:HGNC Symbol;Acc:48349] | RPL36A-HNRNPH2 | chrX:100,645,999-100,667,285 |
|  |  |  |  |  |  |  |  |  | ENSG00000126945 | heterogeneous nuclear ribonucleoprotein H2 (H') [Source:HGNC Symbol;Acc:5042] | HNRNPH2 | chrX:100,663,283-100,669,121 |
|  |  | chrX:118,699,347-118,699,412 | 4 | -1.22 | 7.83e-22 | ENSR00000248346 | Promoter | chrX:118,698,200-118,699,801 | ENSG00000018610 | chromosome X open reading frame 56 [Source:HGNC Symbol;Acc:26239] | CXorf56 | chrX:118,672,112-118,699,397 |
|  |  | chrX:152,989,492-152,990,345 | 18 | -0.10 | 5.65e-05 | ENSR00000249590 | Promoter | chrX:152,987,600-152,993,201 | ENSG00000185825 | B-cell receptor-associated protein 31 [Source:HGNC Symbol;Acc:16695] | BCAP31 | chrX:152,965,947-152,990,152 |
|  |  |  |  |  |  |  |  |  | ENSG00000101986 | ATP-binding cassette, sub-family D (ALD), member 1 [Source:HGNC Symbol;Acc:61] | ABCD1 | chrX:152,990,323-153,010,216 |
|  |  | chrX:153,719,071-153,719,169 | 3 | -0.12 | 8.43e-08 | ENSR00000249678 | Promoter | chrX:153,716,801-153,720,001 |  |  |  |  |
|  | Model 2 | chrX: 15,511,530- 15,512,019 | 9 | 0.62 | 1.73e-03 | ENSR00000244760 | Promoter | chrX:15,510,000-15,512,201 | ENSG00000087842 | pirin (iron-binding nuclear protein) [Source:HGNC Symbol;Acc:30048] | PIR | chrX:15,402,921-15,511,687 |
|  |  |  |  |  |  |  |  |  | ENSG00000102010 | BMX non-receptor tyrosine kinase [Source:HGNC Symbol;Acc:1079] | BMX | chrX:15,482,369-15,574,652 |
|  |  |  |  |  |  | ENSR00000422673 | CTCF Binding Site | chrX:15,511,801-15,512,200 | ENSG00000087842 | pirin (iron-binding nuclear protein) [Source:HGNC Symbol;Acc:30048] | PIR | chrX:15,402,921-15,511,687 |
|  |  |  |  |  |  |  |  |  | ENSG00000102010 | BMX non-receptor tyrosine kinase [Source:HGNC Symbol;Acc:1079] | BMX | chrX:15,482,369-15,574,652 |
|  |  | chrX: 25,034,241- 25,034,688 | 4 | -0.59 | 2.34e-17 | ENSR00002096583 | Promoter | chrX:25,029,001-25,034,600 |  |  |  |  |
|  |  | chrX: 31,284,529- 31,285,029 | 5 | -0.18 | 9.78e-07 | ENSR00000245550 | Promoter | chrX:31,281,801-31,285,401 | ENSG00000198947 | dystrophin [Source:HGNC Symbol;Acc:2928] | DMD | chrX:31,115,794-33,357,558 |
|  |  |  |  |  |  | ENSR00002097003 | CTCF Binding Site | chrX:31,285,001-31,285,200 |  |  |  |  |
|  |  | chrX: 47,863,437- 47,863,595 | 5 | -0.12 | 1.33e-09 | ENSR00000423313 | Promoter | chrX:47,861,600-47,864,400 | ENSG00000221994 | zinc finger protein 630 [Source:HGNC Symbol;Acc:28855] | ZNF630 | chrX:47,842,756-47,931,025 |
|  |  | chrX: 48,768,940- 48,769,091 | 3 | -0.26 | 4.19e-07 | ENSR00000246526 | Promoter | chrX:48,767,200-48,770,801 | ENSG00000102100 | solute carrier family 35 (UDP-galactose transporter), member A2 [Source:HGNC Symbol;Acc:11022] | SLC35A2 | chrX:48,760,459-48,769,235 |
|  |  | chrX: 48,815,747- 48,815,822 | 3 | -0.14 | 5.01e-04 | ENSR00000423354 | Promoter | chrX:48,812,800-48,817,201 |  |  |  |  |
|  |  | chrX: 48,931,572- 48,931,623 | 3 | -0.11 | 5.24e-07 | ENSR00000423364 | Promoter | chrX:48,929,201-48,932,401 | ENSG00000243279 | PRA1 domain family, member 2 [Source:HGNC Symbol;Acc:28911] | PRAF2 | chrX:48,928,813-48,931,730 |
|  |  |  |  |  |  |  |  |  | ENSG00000250232 | WD repeat domain phosphoinositide-interacting protein 4 [Source:UniProtKB/TrEMBL;Acc:A6NM71] | AF196779.12 | chrX:48,929,385-48,937,546 |
|  |  |  |  |  |  |  |  |  | ENSG00000196998 | WD repeat domain 45 [Source:HGNC Symbol;Acc:28912] | WDR45 | chrX:48,929,385-48,958,108 |
|  |  | chrX: 48,931,699- 48,931,851 | 5 | -0.38 | 2.60e-06 |  |  |  | ENSG00000243279 | PRA1 domain family, member 2 [Source:HGNC Symbol;Acc:28911] | PRAF2 | chrX:48,928,813-48,931,730 |
|  |  |  |  |  |  |  |  |  | ENSG00000250232 | WD repeat domain phosphoinositide-interacting protein 4 [Source:UniProtKB/TrEMBL;Acc:A6NM71] | AF196779.12 | chrX:48,929,385-48,937,546 |
|  |  |  |  |  |  |  |  |  | ENSG00000196998 | WD repeat domain 45 [Source:HGNC Symbol;Acc:28912] | WDR45 | chrX:48,929,385-48,958,108 |
|  |  | chrX:100,645,891-100,645,938 | 4 | -0.24 | 3.63e-03 | ENSR00000423844 | Promoter | chrX:100,645,001-100,648,401 | ENSG00000241343 | ribosomal protein L36a [Source:HGNC Symbol;Acc:10359] | RPL36A | chrX:100,645,812-100,651,105 |
|  |  | chrX:100,663,146-100,663,470 | 6 | -0.28 | 3.57e-09 | ENSR00000423848 | Promoter | chrX:100,660,200-100,666,201 | ENSG00000257529 | RPL36A-HNRNPH2 readthrough [Source:HGNC Symbol;Acc:48349] | RPL36A-HNRNPH2 | chrX:100,645,999-100,667,285 |
|  |  |  |  |  |  |  |  |  | ENSG00000126945 | heterogeneous nuclear ribonucleoprotein H2 (H') [Source:HGNC Symbol;Acc:5042] | HNRNPH2 | chrX:100,663,283-100,669,121 |
|  |  | chrX:118,699,347-118,699,475 | 6 | -1.14 | 1.67e-20 | ENSR00000248346 | Promoter | chrX:118,698,200-118,699,801 | ENSG00000018610 | chromosome X open reading frame 56 [Source:HGNC Symbol;Acc:26239] | CXorf56 | chrX:118,672,112-118,699,397 |
|  |  | chrX:152,989,492-152,990,341 | 17 | -0.11 | 3.73e-05 | ENSR00000249590 | Promoter | chrX:152,987,600-152,993,201 | ENSG00000185825 | B-cell receptor-associated protein 31 [Source:HGNC Symbol;Acc:16695] | BCAP31 | chrX:152,965,947-152,990,152 |
|  |  |  |  |  |  |  |  |  | ENSG00000101986 | ATP-binding cassette, sub-family D (ALD), member 1 [Source:HGNC Symbol;Acc:61] | ABCD1 | chrX:152,990,323-153,010,216 |
|  |  | chrX:153,719,071-153,719,169 | 3 | -0.11 | 7.60e-06 | ENSR00000249678 | Promoter | chrX:153,716,801-153,720,001 |  |  |  |  |
|  | Model 3 | chrX: 15,511,530- 15,512,019 | 9 | 0.60 | 3.11e-03 | ENSR00000244760 | Promoter | chrX:15,510,000-15,512,201 | ENSG00000087842 | pirin (iron-binding nuclear protein) [Source:HGNC Symbol;Acc:30048] | PIR | chrX:15,402,921-15,511,687 |
|  |  |  |  |  |  |  |  |  | ENSG00000102010 | BMX non-receptor tyrosine kinase [Source:HGNC Symbol;Acc:1079] | BMX | chrX:15,482,369-15,574,652 |
|  |  |  |  |  |  | ENSR00000422673 | CTCF Binding Site | chrX:15,511,801-15,512,200 | ENSG00000087842 | pirin (iron-binding nuclear protein) [Source:HGNC Symbol;Acc:30048] | PIR | chrX:15,402,921-15,511,687 |
|  |  |  |  |  |  |  |  |  | ENSG00000102010 | BMX non-receptor tyrosine kinase [Source:HGNC Symbol;Acc:1079] | BMX | chrX:15,482,369-15,574,652 |
|  |  | chrX: 25,034,241- 25,034,688 | 4 | -0.58 | 1.04e-17 | ENSR00002096583 | Promoter | chrX:25,029,001-25,034,600 |  |  |  |  |
|  |  | chrX: 31,284,529- 31,285,029 | 5 | -0.18 | 2.39e-06 | ENSR00000245550 | Promoter | chrX:31,281,801-31,285,401 | ENSG00000198947 | dystrophin [Source:HGNC Symbol;Acc:2928] | DMD | chrX:31,115,794-33,357,558 |
|  |  |  |  |  |  | ENSR00002097003 | CTCF Binding Site | chrX:31,285,001-31,285,200 |  |  |  |  |
|  |  | chrX: 47,863,437- 47,863,595 | 5 | -0.13 | 3.35e-13 | ENSR00000423313 | Promoter | chrX:47,861,600-47,864,400 | ENSG00000221994 | zinc finger protein 630 [Source:HGNC Symbol;Acc:28855] | ZNF630 | chrX:47,842,756-47,931,025 |
|  |  | chrX: 48,768,940- 48,769,091 | 3 | -0.24 | 7.11e-06 | ENSR00000246526 | Promoter | chrX:48,767,200-48,770,801 | ENSG00000102100 | solute carrier family 35 (UDP-galactose transporter), member A2 [Source:HGNC Symbol;Acc:11022] | SLC35A2 | chrX:48,760,459-48,769,235 |
|  |  | chrX: 48,815,747- 48,815,822 | 3 | -0.12 | 4.83e-03 | ENSR00000423354 | Promoter | chrX:48,812,800-48,817,201 |  |  |  |  |
|  |  | chrX: 48,931,572- 48,931,623 | 3 | -0.11 | 2.59e-06 | ENSR00000423364 | Promoter | chrX:48,929,201-48,932,401 | ENSG00000243279 | PRA1 domain family, member 2 [Source:HGNC Symbol;Acc:28911] | PRAF2 | chrX:48,928,813-48,931,730 |
|  |  |  |  |  |  |  |  |  | ENSG00000250232 | WD repeat domain phosphoinositide-interacting protein 4 [Source:UniProtKB/TrEMBL;Acc:A6NM71] | AF196779.12 | chrX:48,929,385-48,937,546 |
|  |  |  |  |  |  |  |  |  | ENSG00000196998 | WD repeat domain 45 [Source:HGNC Symbol;Acc:28912] | WDR45 | chrX:48,929,385-48,958,108 |
|  |  | chrX: 48,931,699- 48,931,851 | 5 | -0.38 | 4.67e-06 |  |  |  | ENSG00000243279 | PRA1 domain family, member 2 [Source:HGNC Symbol;Acc:28911] | PRAF2 | chrX:48,928,813-48,931,730 |
|  |  |  |  |  |  |  |  |  | ENSG00000250232 | WD repeat domain phosphoinositide-interacting protein 4 [Source:UniProtKB/TrEMBL;Acc:A6NM71] | AF196779.12 | chrX:48,929,385-48,937,546 |
|  |  |  |  |  |  |  |  |  | ENSG00000196998 | WD repeat domain 45 [Source:HGNC Symbol;Acc:28912] | WDR45 | chrX:48,929,385-48,958,108 |
|  |  | chrX:100,663,213-100,663,543 | 3 | -0.50 | 2.82e-06 | ENSR00000423848 | Promoter | chrX:100,660,200-100,666,201 | ENSG00000257529 | RPL36A-HNRNPH2 readthrough [Source:HGNC Symbol;Acc:48349] | RPL36A-HNRNPH2 | chrX:100,645,999-100,667,285 |
|  |  |  |  |  |  |  |  |  | ENSG00000126945 | heterogeneous nuclear ribonucleoprotein H2 (H') [Source:HGNC Symbol;Acc:5042] | HNRNPH2 | chrX:100,663,283-100,669,121 |
|  |  | chrX:118,699,347-118,699,412 | 4 | -1.25 | 1.09e-22 | ENSR00000248346 | Promoter | chrX:118,698,200-118,699,801 | ENSG00000018610 | chromosome X open reading frame 56 [Source:HGNC Symbol;Acc:26239] | CXorf56 | chrX:118,672,112-118,699,397 |
|  |  | chrX:152,989,492-152,990,345 | 18 | -0.10 | 8.02e-05 | ENSR00000249590 | Promoter | chrX:152,987,600-152,993,201 | ENSG00000185825 | B-cell receptor-associated protein 31 [Source:HGNC Symbol;Acc:16695] | BCAP31 | chrX:152,965,947-152,990,152 |
|  |  |  |  |  |  |  |  |  | ENSG00000101986 | ATP-binding cassette, sub-family D (ALD), member 1 [Source:HGNC Symbol;Acc:61] | ABCD1 | chrX:152,990,323-153,010,216 |
|  |  | chrX:153,719,071-153,719,169 | 3 | -0.13 | 2.06e-08 | ENSR00000249678 | Promoter | chrX:153,716,801-153,720,001 |  |  |  |  |
|  | Model 4 | chrX: 15,511,530- 15,512,019 | 9 | 0.63 | 8.09e-04 | ENSR00000244760 | Promoter | chrX:15,510,000-15,512,201 | ENSG00000087842 | pirin (iron-binding nuclear protein) [Source:HGNC Symbol;Acc:30048] | PIR | chrX:15,402,921-15,511,687 |
|  |  |  |  |  |  |  |  |  | ENSG00000102010 | BMX non-receptor tyrosine kinase [Source:HGNC Symbol;Acc:1079] | BMX | chrX:15,482,369-15,574,652 |
|  |  |  |  |  |  | ENSR00000422673 | CTCF Binding Site | chrX:15,511,801-15,512,200 | ENSG00000087842 | pirin (iron-binding nuclear protein) [Source:HGNC Symbol;Acc:30048] | PIR | chrX:15,402,921-15,511,687 |
|  |  |  |  |  |  |  |  |  | ENSG00000102010 | BMX non-receptor tyrosine kinase [Source:HGNC Symbol;Acc:1079] | BMX | chrX:15,482,369-15,574,652 |
|  |  | chrX: 25,034,241- 25,034,688 | 4 | -0.59 | 5.26e-18 | ENSR00002096583 | Promoter | chrX:25,029,001-25,034,600 |  |  |  |  |
|  |  | chrX: 31,284,529- 31,285,029 | 5 | -0.18 | 2.94e-06 | ENSR00000245550 | Promoter | chrX:31,281,801-31,285,401 | ENSG00000198947 | dystrophin [Source:HGNC Symbol;Acc:2928] | DMD | chrX:31,115,794-33,357,558 |
|  |  |  |  |  |  | ENSR00002097003 | CTCF Binding Site | chrX:31,285,001-31,285,200 |  |  |  |  |
|  |  | chrX: 47,863,437- 47,863,529 | 3 | -0.13 | 3.61e-10 | ENSR00000423313 | Promoter | chrX:47,861,600-47,864,400 | ENSG00000221994 | zinc finger protein 630 [Source:HGNC Symbol;Acc:28855] | ZNF630 | chrX:47,842,756-47,931,025 |
|  |  | chrX: 48,768,940- 48,769,091 | 3 | -0.25 | 9.80e-07 | ENSR00000246526 | Promoter | chrX:48,767,200-48,770,801 | ENSG00000102100 | solute carrier family 35 (UDP-galactose transporter), member A2 [Source:HGNC Symbol;Acc:11022] | SLC35A2 | chrX:48,760,459-48,769,235 |
|  |  | chrX: 48,815,747- 48,815,822 | 3 | -0.13 | 1.73e-03 | ENSR00000423354 | Promoter | chrX:48,812,800-48,817,201 |  |  |  |  |
|  |  | chrX: 48,931,572- 48,931,623 | 3 | -0.12 | 1.64e-07 | ENSR00000423364 | Promoter | chrX:48,929,201-48,932,401 | ENSG00000243279 | PRA1 domain family, member 2 [Source:HGNC Symbol;Acc:28911] | PRAF2 | chrX:48,928,813-48,931,730 |
|  |  |  |  |  |  |  |  |  | ENSG00000250232 | WD repeat domain phosphoinositide-interacting protein 4 [Source:UniProtKB/TrEMBL;Acc:A6NM71] | AF196779.12 | chrX:48,929,385-48,937,546 |
|  |  |  |  |  |  |  |  |  | ENSG00000196998 | WD repeat domain 45 [Source:HGNC Symbol;Acc:28912] | WDR45 | chrX:48,929,385-48,958,108 |
|  |  | chrX: 48,931,699- 48,931,851 | 5 | -0.37 | 8.67e-06 |  |  |  | ENSG00000243279 | PRA1 domain family, member 2 [Source:HGNC Symbol;Acc:28911] | PRAF2 | chrX:48,928,813-48,931,730 |
|  |  |  |  |  |  |  |  |  | ENSG00000250232 | WD repeat domain phosphoinositide-interacting protein 4 [Source:UniProtKB/TrEMBL;Acc:A6NM71] | AF196779.12 | chrX:48,929,385-48,937,546 |
|  |  |  |  |  |  |  |  |  | ENSG00000196998 | WD repeat domain 45 [Source:HGNC Symbol;Acc:28912] | WDR45 | chrX:48,929,385-48,958,108 |
|  |  | chrX:100,645,891-100,645,938 | 4 | -0.25 | 2.45e-03 | ENSR00000423844 | Promoter | chrX:100,645,001-100,648,401 | ENSG00000241343 | ribosomal protein L36a [Source:HGNC Symbol;Acc:10359] | RPL36A | chrX:100,645,812-100,651,105 |
|  |  | chrX:100,663,146-100,663,213 | 5 | -0.29 | 4.27e-09 | ENSR00000423848 | Promoter | chrX:100,660,200-100,666,201 | ENSG00000257529 | RPL36A-HNRNPH2 readthrough [Source:HGNC Symbol;Acc:48349] | RPL36A-HNRNPH2 | chrX:100,645,999-100,667,285 |
|  |  | chrX:118,699,347-118,699,475 | 6 | -1.16 | 1.77e-21 | ENSR00000248346 | Promoter | chrX:118,698,200-118,699,801 | ENSG00000018610 | chromosome X open reading frame 56 [Source:HGNC Symbol;Acc:26239] | CXorf56 | chrX:118,672,112-118,699,397 |
|  |  | chrX:153,060,015-153,060,147 | 3 | 0.24 | 4.49e-04 | ENSR00000249598 | Promoter | chrX:153,058,200-153,062,201 | ENSG00000180879 | signal sequence receptor, delta [Source:HGNC Symbol;Acc:11326] | SSR4 | chrX:153,058,971-153,063,960 |
|  |  | chrX:153,719,071-153,719,169 | 3 | -0.11 | 2.97e-06 | ENSR00000249678 | Promoter | chrX:153,716,801-153,720,001 |  |  |  |  |
