## Supplementary figures and tables for "The X-factor in ART: does the use of Assisted Reproductive Technologies influence DNA methylation on the X chromosome?"

September 22, 2022

**Affiliations:**
[1] Centre for Fertility and Health, Norwegian Institute of Public Health, 0213 Oslo, Norway
[2] Department of Global Public Health and Primary Care, University of Bergen, 5020 Bergen, Norway
[3] Deepinsight, 0154 Oslo, Norway
[4] Department of Mathematics, Faculty of Mathematics and Natural Sciences, University of Oslo, 0315 Oslo, Norway
[5] Department of Human Genetics, University of Chicago, Chicago, IL 60637, USA
[6] Department of Method Development and Analytics, Norwegian Institute of Public Health, 0213 Oslo, Norway
[7] Department of Computer Science, Electrical Engineering and Mathematical Sciences, Western Norway University of Applied Sciences, 5020 Bergen, Norway
[8] Murdoch Children's Research Institute, Melbourne, Victoria 3052, Australia
[9] Department of Paediatrics, University of Melbourne, Victoria 3010, Australia
[10] Department of Medical Genetics, Oslo University Hospital and University of Oslo, 0424 Oslo, Norway

**Corresponding author:*
Julia Romanowska, PhD
Department of Global Public Health and Primary Care
University of Bergen
5020 Bergen, Norway


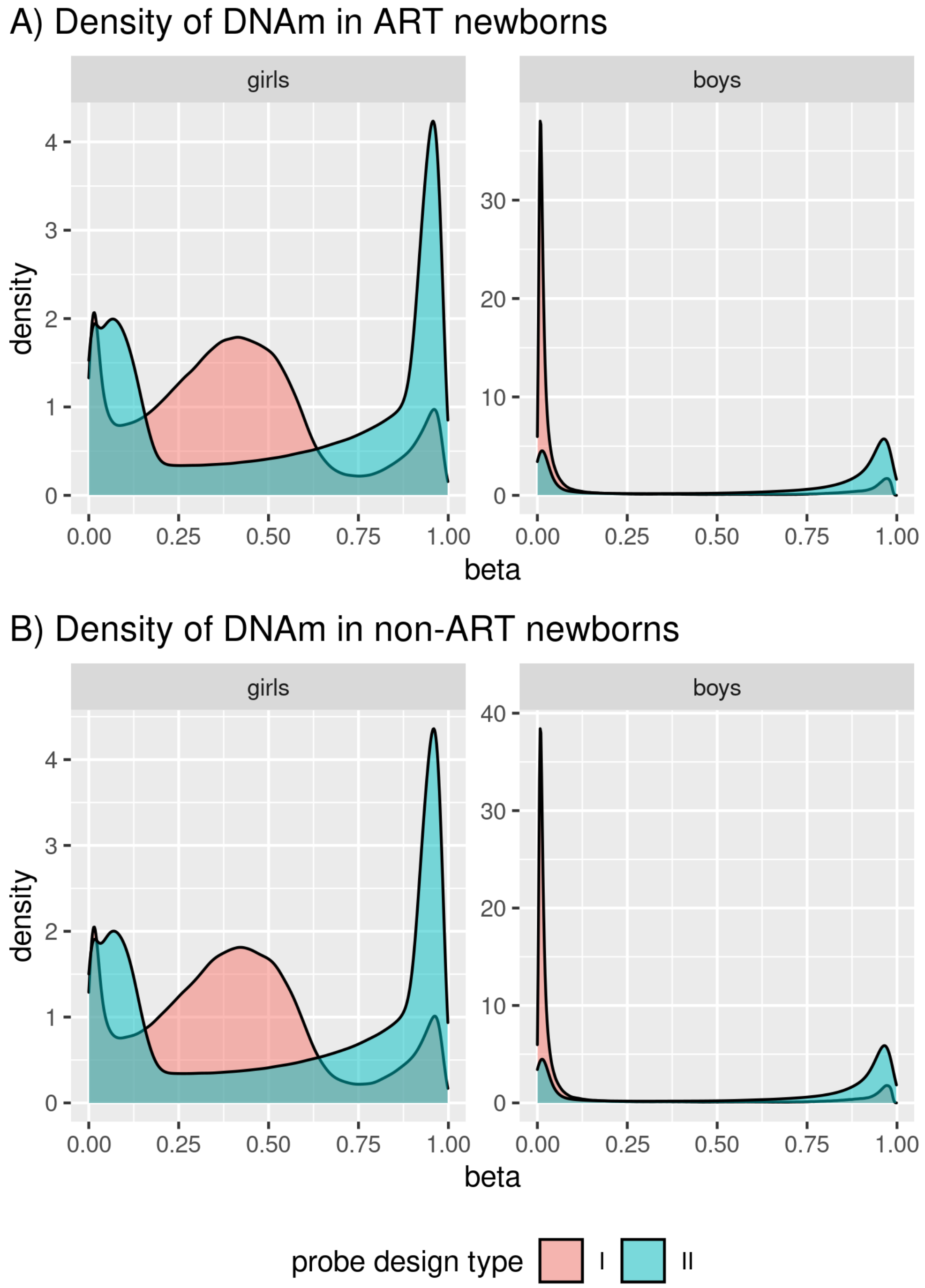

Figure S1: Sex-stratified density plots of the X-chromosome $\beta$-values for ART and non-ART children according to Type I and Type II probes on the Illumina EPIC array.

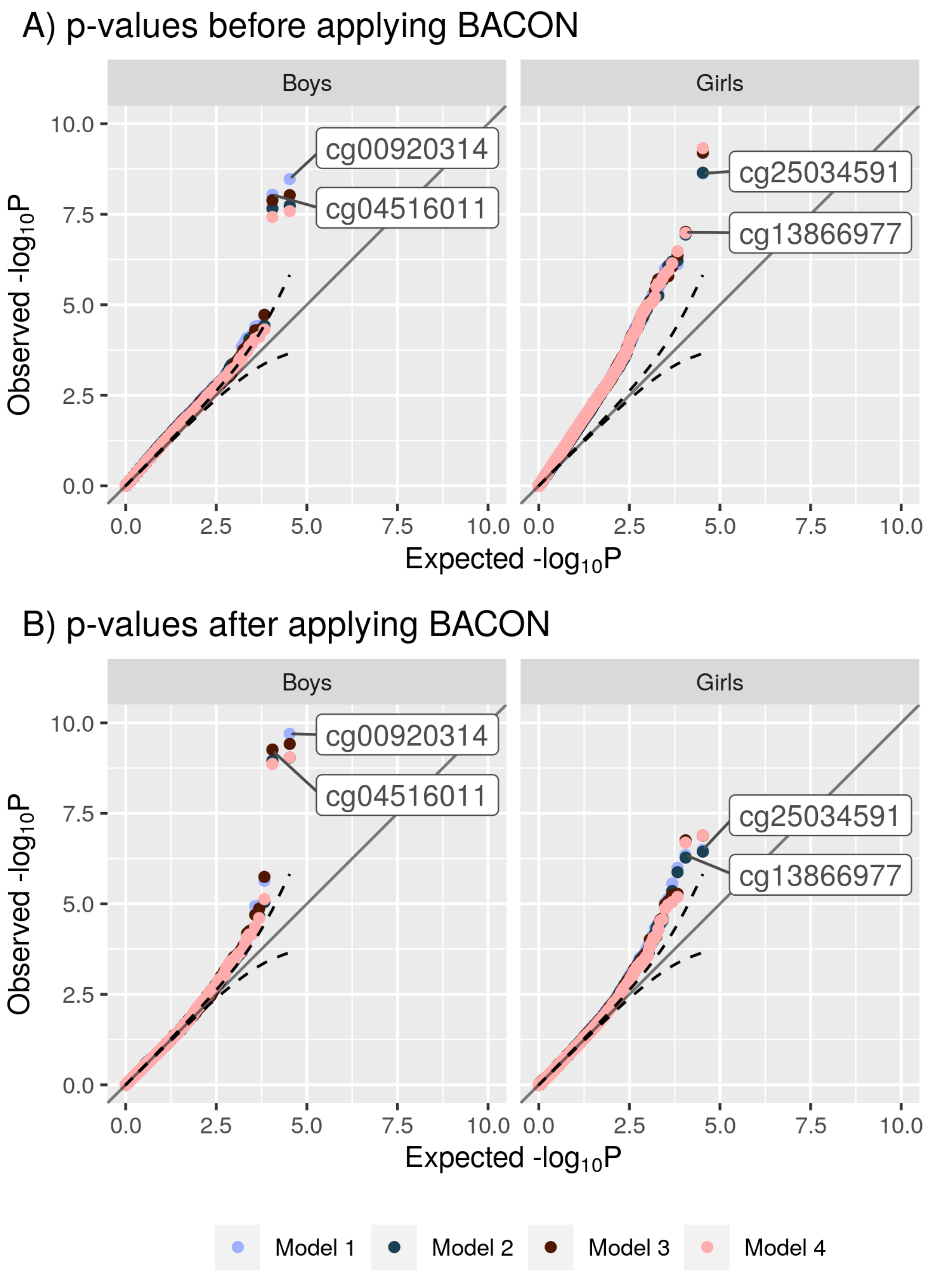

Figure S2: Quantile-quantile (QQ) plot of the observed versus expected $-\log_{10} p$-values of the results for boys and girls before (panel A) and after (panel B) adjustment with BACON.

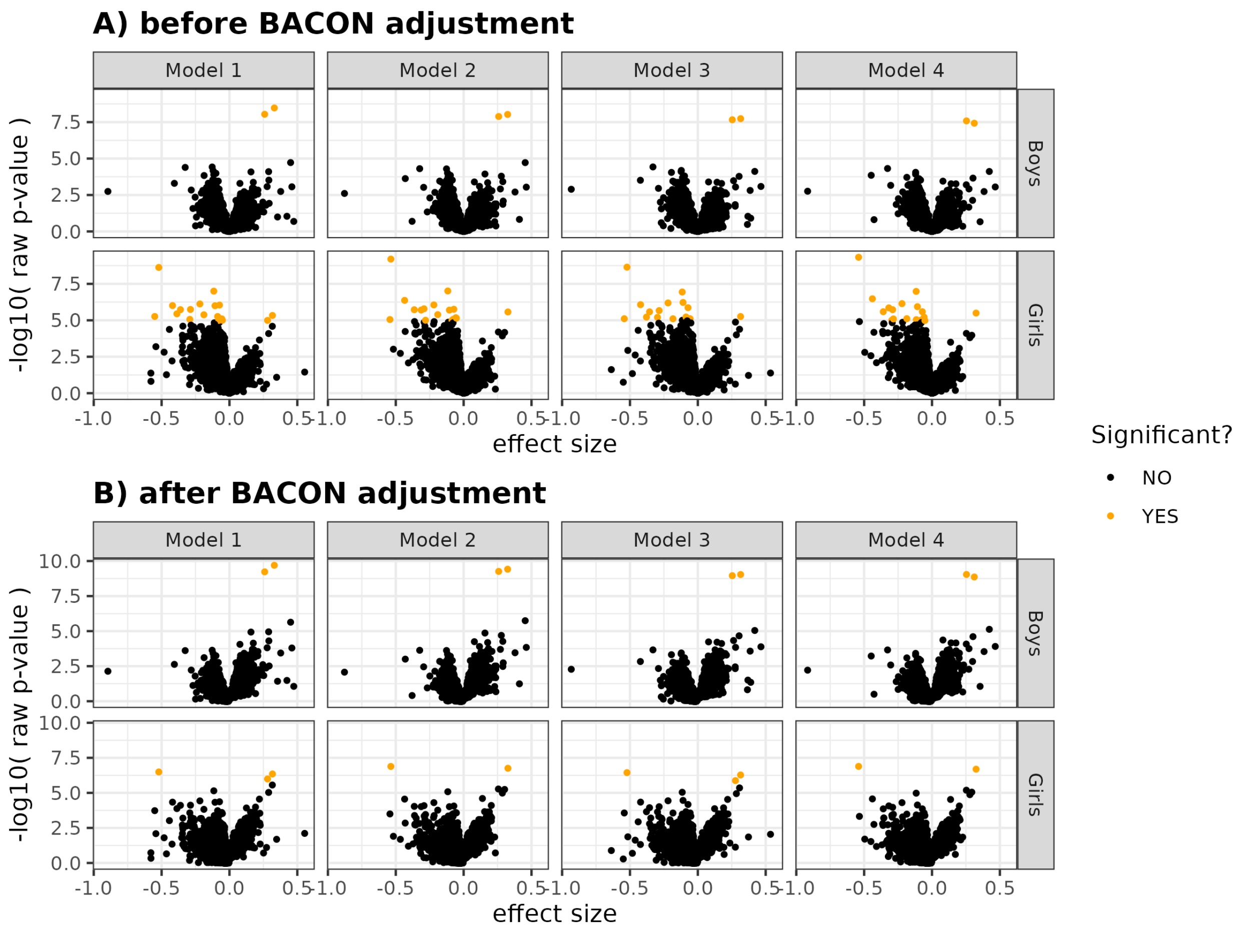

Figure S3: Effect sizes versus $-\log_{10} p$-values for each of the X-linked CpGs included in the analyses. Significant findings at FDR $< 0.05$ are highlighted in orange.

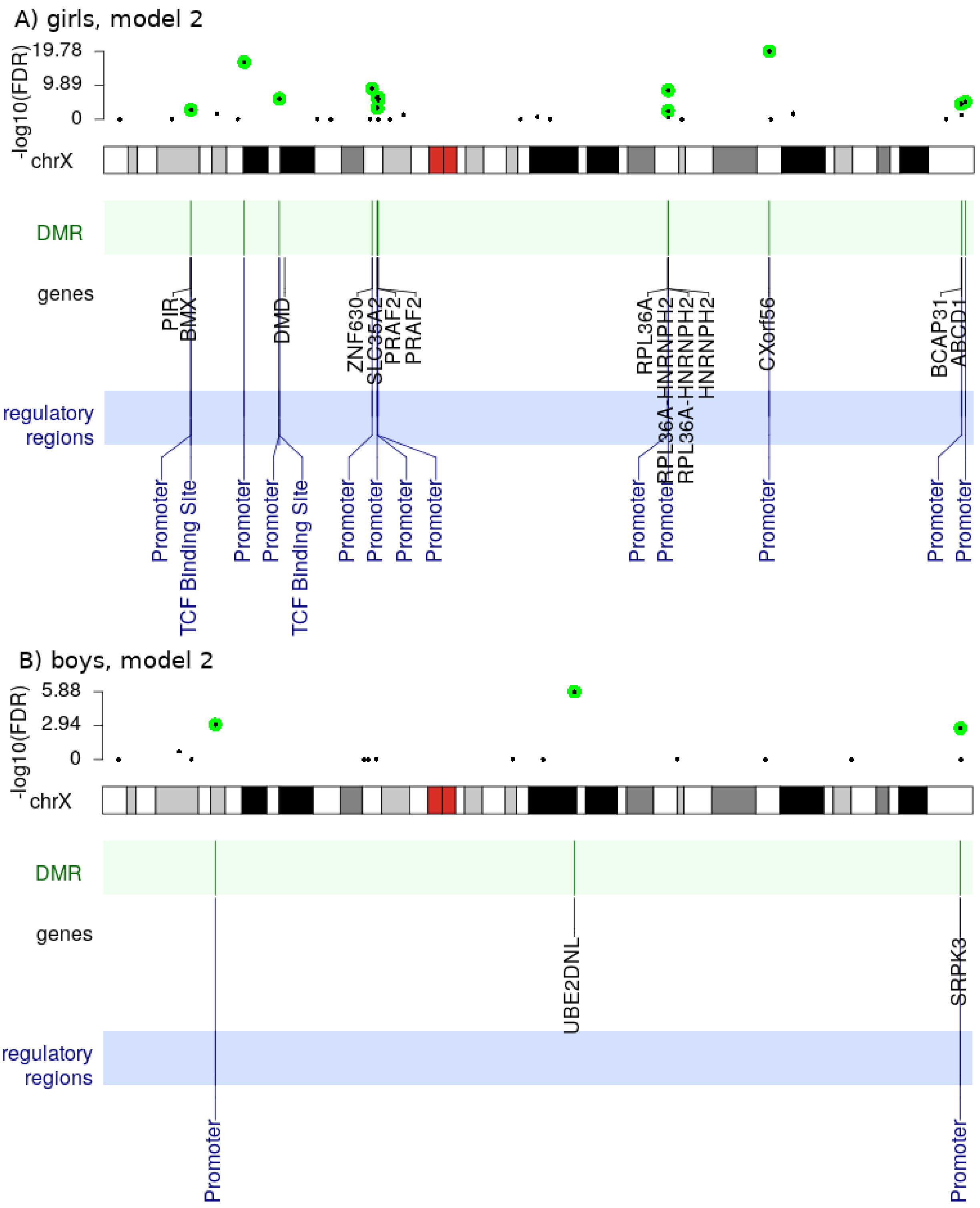

Figure S4: Location of DMRs on the X chromosome for the analyses of Model 2. The top part of each panel shows p-values for all the DMRs that contained at least three CpGs and the FDR-adjusted p-values $< 0.01$ are marked green. The bottom part of each figure marks genes and regulatory regions harbored by the significant DMRs.

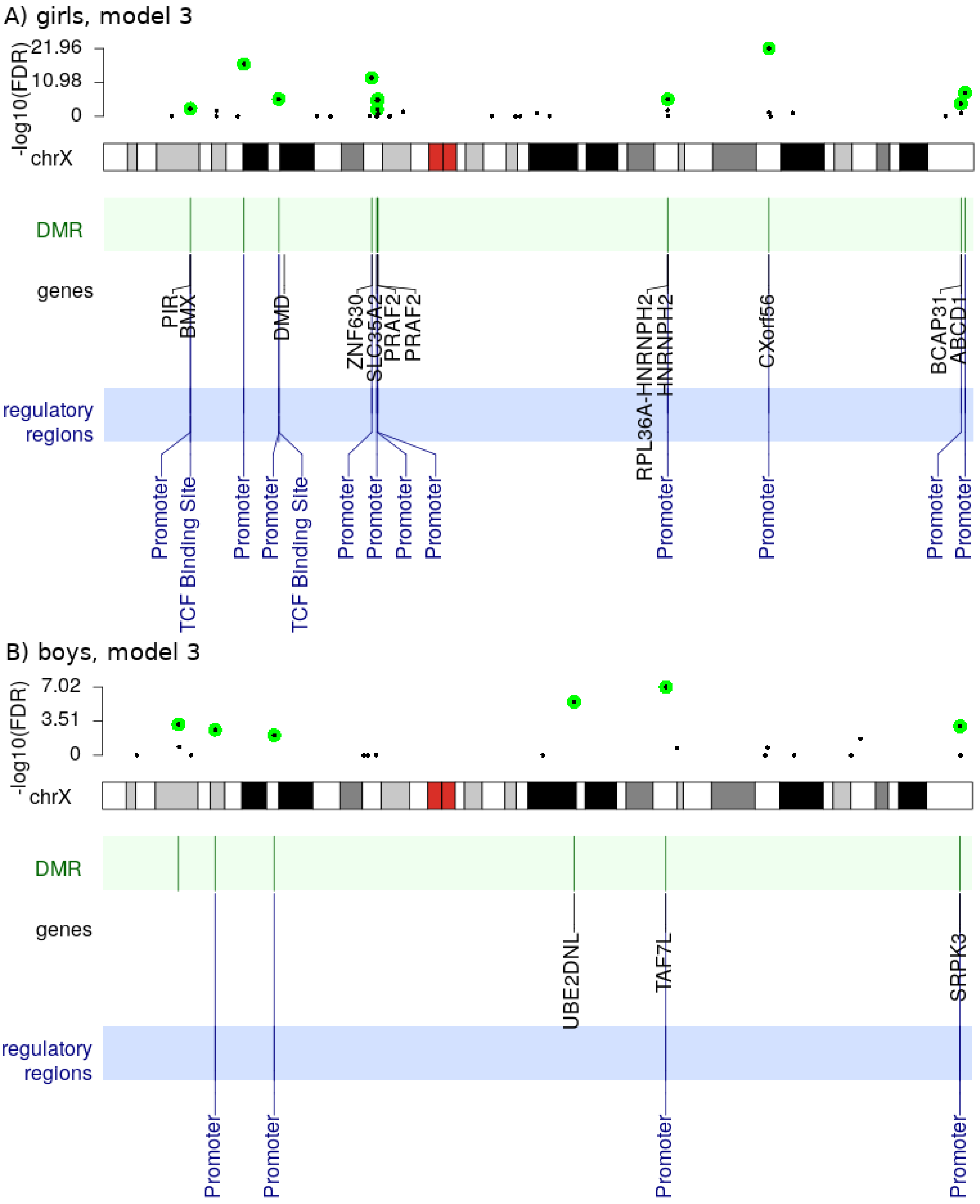

Figure S5: Location of DMRs on the X chromosome for the analyses of Model 3. The top part of each panel shows p-values for all the DMRs that contained at least three CpGs, and the FDR-adjusted p-values $< 0.01$ are marked green. The bottom part of each figure marks genes and regulatory regions harbored by the significant DMRs.

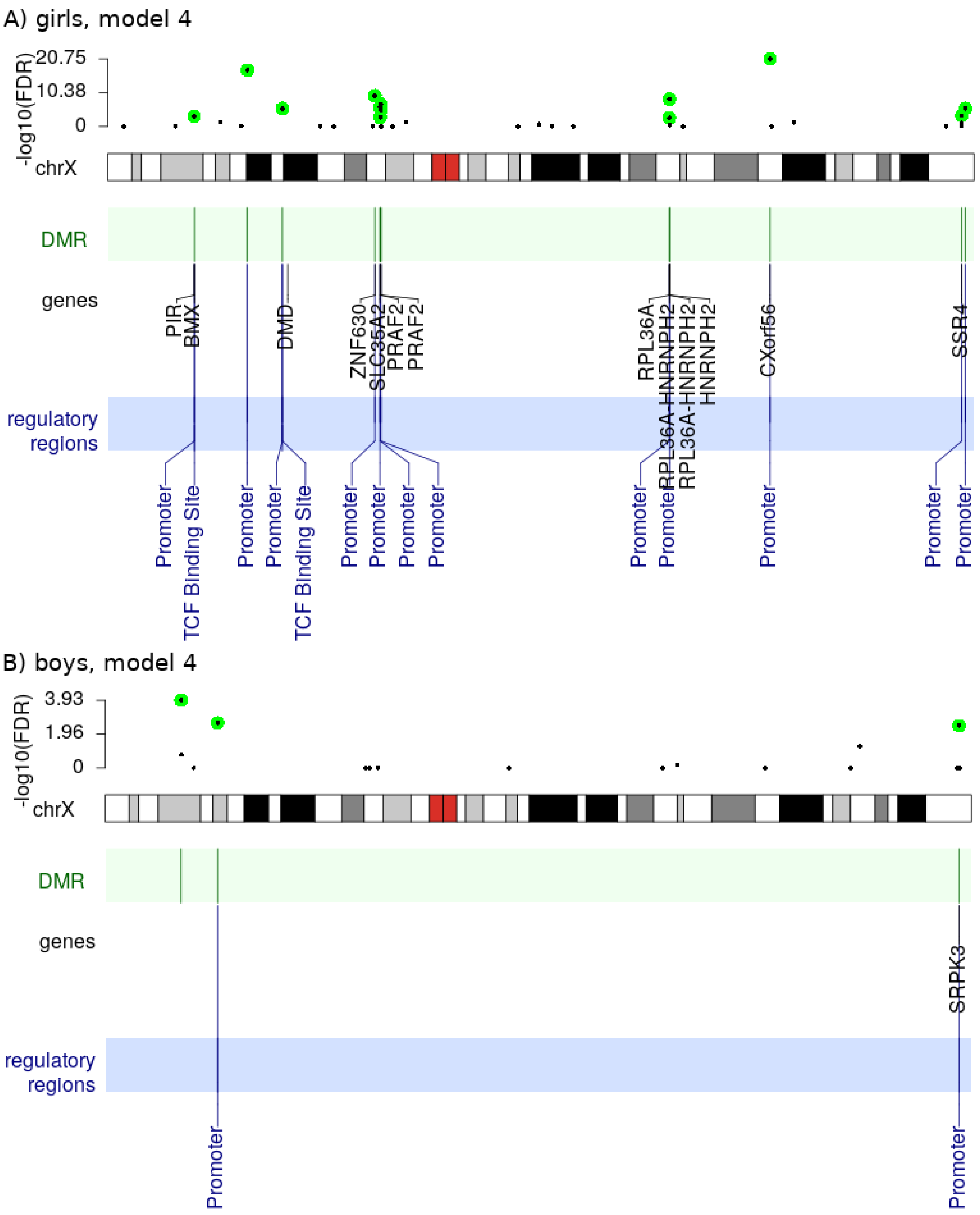

Figure S6: Location of DMRs on the X chromosome for the analyses of Model 4. The top part of each panel shows p-values for all the DMRs that contained at least three CpGs, and the FDR-adjusted p-values $< 0.01$ are marked green. The bottom part of each figure marks genes and regulatory regions harbored by the significant DMRs.

Figure S7: DNA methylation at two CpGs within the *AMOT* and *EIF2S3* genes in CHART dataset. Note that these two genes were among the significant findings in the girls-only XWAS if the MoBa dataset. Abbreviations: F=female, M=male, Ctrl=non-ART newborn, ART=newborn conceived through the use of ART.

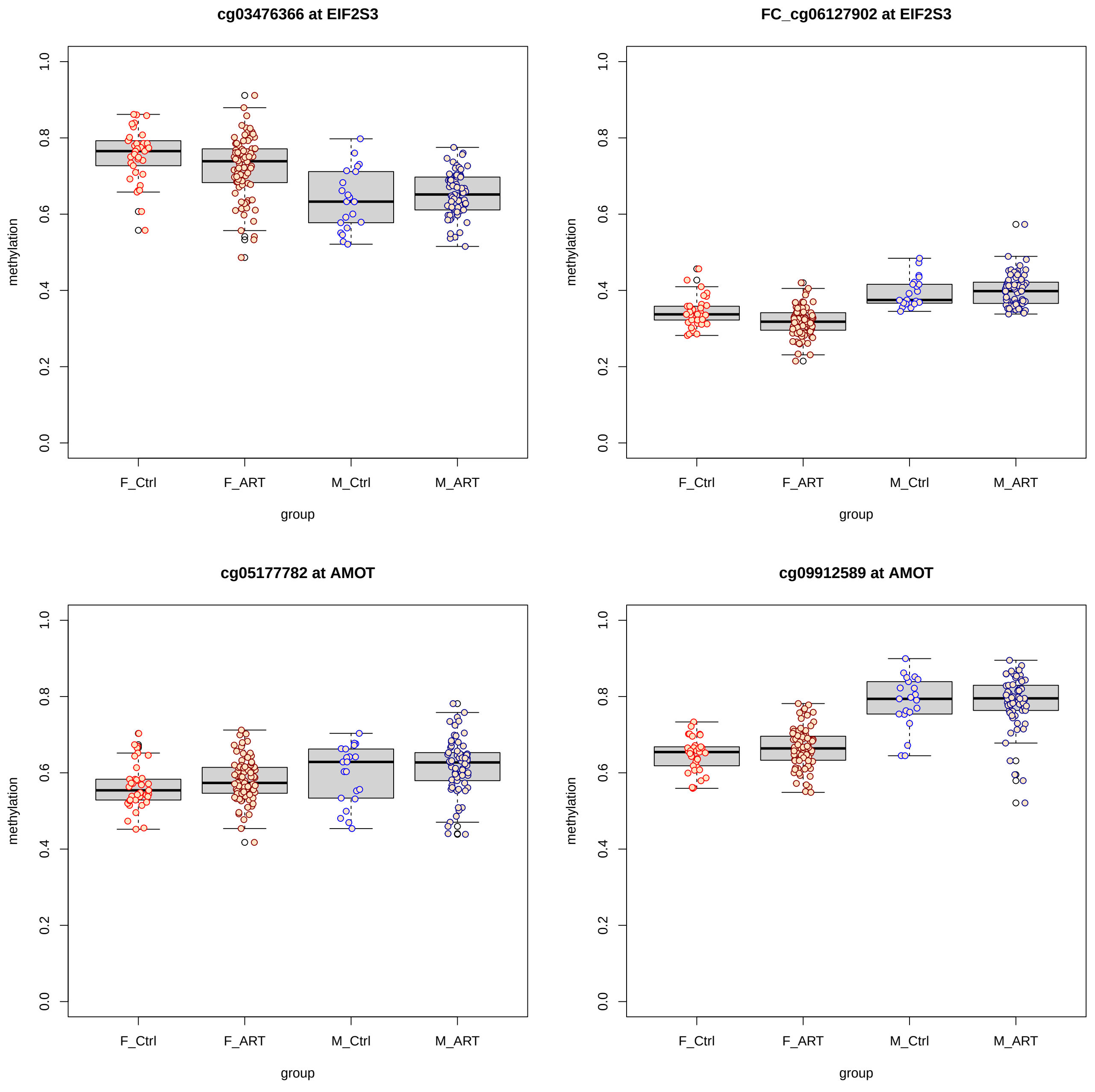

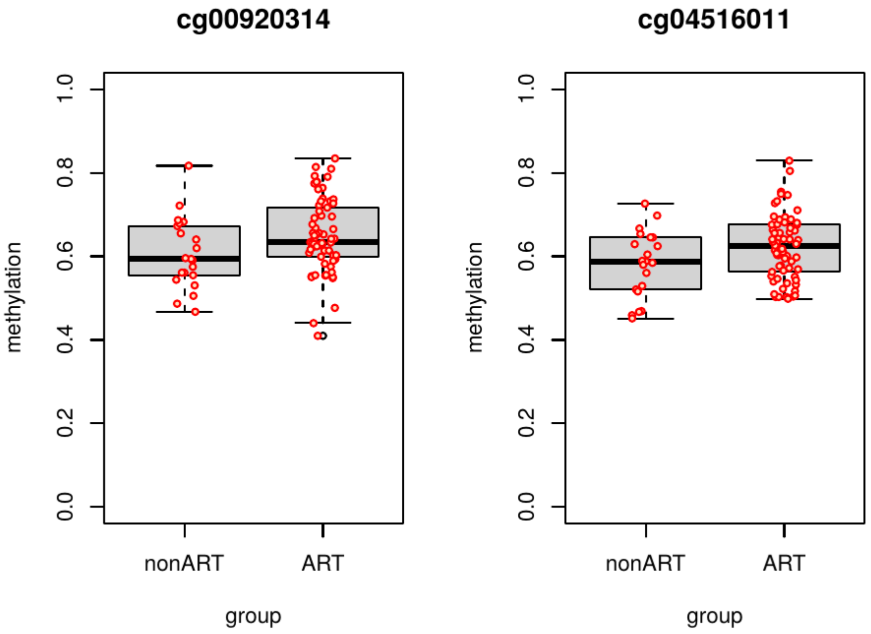

Figure S8: DNA methylation at two CpGs within the *UBE2DNL* pseudogene in the CHART dataset. Note that these two CpGs were significant and robust findings in the boys-only XWAS of the MoBa data.

Table S1: Genes found to be possibly regulated by regulatory regions that co-localized with the most significant XWAS findings.

| Gene name | Product[(a)] | Function | Expressed in[(b)] | Refs and URLs |
|---|---|---|---|---|
| | | *ensembl regulatory ID ENSR00000245352 (co-localized with cg25034591)* | | |
| *NONHSAG054159.2* | lncRNA | Putative role in transcription regulation | Not tissue specific (https://ngdc.cncb.ac.cn/lncbook/gene?geneid=HSALNG0137031) | Statello, L., et al. Nat Rev Mol Cell Biol 22, 96–118 (2021). https://doi.org/10.1038/s41580-020-00315-9 |
| *RF00017-7804* | SRP_RNA | Putative role in transcription regulation | No data | Akopian, D., et al. Annu Rev Biochem. 2013; 82: 693–721. https://doi.org/10.1146/annurev-biochem-072711-164732 |
| *RF00017-7805* | SRP_RNA | Putative role in transcription regulation | No data | Akopian, D., et al. Annu Rev Biochem. 2013 ; 82: 693–721. https://doi.org/10.1146/annurev-biochem-072711-164732 |
| *HSALNG0137030* | lncRNA | Putative role in transcription regulation | Not tissue specific (https://ngdc.cncb.ac.cn/lncbook/gene?geneid=HSALNG0137030) | Statello, L., et al. Nat Rev Mol Cell Biol 22, 96–118 (2021). https://doi.org/10.1038/s41580-020-00315-9 |
| *RF00017-7813* | SRP_RNA | Putative role in transcription regulation | No data | Akopian, D., et al. Annu Rev Biochem. 2013 ; 82: 693–721. https://doi.org/10.1146/annurev-biochem-072711-164732 |
| *EIF2S3* | Eukaryotic Translation Initiation Factor 2 Subunit Gamma | translation initiation | Not tissue specific (https://www.proteinatlas.org/ENSG00000130741-EIF2S3/tissue) | https://www.uniprot.org/uniprot/P41091 |
| *POLA1* | DNA Polymerase Alpha 1, Catalytic Subunit | controls DNA repair and replication | Not tissue specific (https://www.proteinatlas.org/ENSG00000101868-POLA1/tissue) | https://www.uniprot.org/uniprot/P09884 |
| *ZFX* | Zinc Finger Protein X-Linked | transcriptional activator during processes such as oocyte development and spermatogenesis | Not tissue specific (https://www.proteinatlas.org/ENSG00000005889-ZFX/tissue) | https://www.uniprot.org/uniprot/P17010 |
| *KLHL15* | Kelch Like Family Member 15 | protein ubiquitination | Tissue enhanced (bone marrow) (https://www.proteinatlas.org/ENSG00000174010-KLHL15/tissue) | https://www.uniprot.org/uniprot/Q96M94 |
| *RPL9* | Ribosomal Protein L9 Pseudogene 7 | ribosomal protein, thus involved in translation | Not tissue specific (https://www.proteinatlas.org/ENSG00000163682-RPL9/tissue) | https://www.uniprot.org/uniprot/P32969 |
| | | *ensembl regulatory ID ENSR00000912938 (a.k.a. ENSR00001768065) (co-localized with cg13866977)* | | |
| *AMOT* | Angiomotin | Important role during formation of new blood vessels in placenta | Tissue enhanced (epididymis/testis, tongue) (https://www.proteinatlas.org/ENSG00000126016-AMOT/tissue) | https://www.uniprot.org/uniprot/Q4VCS5 |
| *LHFPL1* | LHFPL Tetraspan Subfamily Member 1 | Transmembrane protein | Group enriched (brain, salivary gland) (https://www.proteinatlas.org/ENSG00000182508-LHFPL1/tissue) | https://www.uniprot.org/uniprot/Q86WI0 |
| *MIR4329* | miRNA | Post-transcriptional regulation of gene expression | No data | https://www.nature.com/articles/s41576-020-0263-7 https://www.nature.com/articles/s41576-020-00309-5 |
| *piR-39314* | piRNA | cleaving tRNA, promoting heterochromatin assembly and methylating DNA | No data | Ozata, D.M., A. et al. Nat Rev Genet 20, 89–108 (2019) https://doi.org/10.1038/s41576-018-0073-3 |

[(a)] lncRNA = long non-coding RNA; SRP_RNA = signal recognition particle RNA; miRNA = micro RNA; piRNA = Piwi-interacting RNA;
[(b)] n.a. = not applicable

Table S2: Genes found possibly regulated by regulatory region colocalized with the most significant differentially methylated regions (DMRs).

| Gene name | Product[a] | Function | Expressed in[b] | Refs and URLs |
|---|---|---|---|---|
| | | *ensembl ID ENSR00000248346 (co-localized with DMR chrX:118,699,347-118,699,412)* | | |
| *STEEP1* (aka *CXorf56*) | STING1 ER Exit Protein 1 | positive regulator of STING signaling, an oligomer essential for proper immune response | No data | https://www.uniprot.org/uniprot/Q9H5V9; Zhang, B., Nature Immunology (2020) https://doi.org/10.1038/s41590-020-0730-5 |
| *SLC25A5* | Solute Carrier Family 25 Member 5 | ADP/ATP translocase 2, important in mitochondrial processes [PMID: 31883789] and was also shown to be part of chromosome segregation process [PMID: 20797633]; "Suppressed expression of this gene has been shown to induce apoptosis and inhibit tumor growth". (via GeneCards) | Not tissue specific (https://www.proteinatlas.org/ENSG00000005022-SLC25A5/tissue) | https://www.uniprot.org/uniprot/P05141; Ito, S., et al. Molecular Cell (2010) https://doi.org/10.1016/j.molcel.2010.07.029; Namba, T., et al., Neuron (2020) https://doi.org/10.1016/j.neuron.2019.11.027 |
| *piR-52079-224* | piRNA | cleaving tRNA, promoting heterochromatin assembly and methylating DNA | No data | Ozata, D.M., A. et al. Nat Rev Genet 20, 89–108 (2019) https://doi.org/10.1038/s41576-018-0073-3 |
| | | *ensembl ID ENSR00000249590 (co-localized with DMR chrX:152,989,492-152,990,345)* | | |
| *SLC6A8* | Solute Carrier Family 6 Member 8 | "transports creatine into and out of cells. Defects in this gene can result in X-linked creatine deficiency syndrome" (via GeneCards) | RNA expression mainly in mitochondria, low tissue specificity (https://www.proteinatlas.org/ENSG00000130821-SLC6A8/tissue) | https://www.uniprot.org/uniprot/P48029 |
| *ABCD1* | ATP Binding Cassette Subfamily D Member 1 | "plays a role in regulation of VLCFAs and energy metabolism", "Controls also the cellular response to oxidative stress by regulating mitochondrial functions [...]. And finally controls the inflammatory response" (via UniProt) | RNA expression mainly in mitochondria, low tissue specificity (https://www.proteinatlas.org/ENSG00000101986-ABCD1/tissue) | https://www.uniprot.org/uniprot/P33897 |
| *BCAP31* | B Cell Receptor Associated Protein 31 | Chaperone protein in endoplasmic reticulum (ER), important in mitochondrial function | Low tissue specificity; in fibroblasts clustered mainly with other genes of hormone signalling pathway (https://www.proteinatlas.org/ENSG00000185825-BCAP31/tissue) | https://www.uniprot.org/uniprot/P51572 |
| *PLXNB3* | Plexin B3 | "plays a role in axon guidance, invasive growth and cell migration" (via GeneCards) | RNA expression enriched in brain and clustered with other genes of myelination pathway (https://www.proteinatlas.org/ENSG00000198753-PLXNB3/tissue) | https://www.uniprot.org/uniprot/Q9ULL4 |
| *PNCK* | Pregnancy Up-Regulated Nonubiquitous CaM Kinase | "Phosphorylates and activates CAMK1" (via UniProt) | RNA expression enriched in brain and clustered with other genes of ion transport pathway (https://www.proteinatlas.org/ENSG00000130822-PNCK/tissue) | https://www.uniprot.org/uniprot/Q6P2M8 |
| *PDZD4* | PDZ Domain Containing 4 | Brain-specific protein; ubiquitin protein ligase activity | RNA expression enriched in brain and clustered with other genes of ion transport pathway (https://www.proteinatlas.org/ENSG00000067840-PDZD4/tissue) | https://www.uniprot.org/uniprot/Q76G19 |
| *KRT18P48* | Keratin 18 Pseudogene 48 | pseudogene | n.a. | n.a. |
| *HSALNG0140788* | lncRNA | Putative role in transcription regulation | Not tissue specific (https://ngdc.cncb.ac.cn/lncbook/gene?geneid=HSALNG0140785) | n.a. |
| *HSALNG0140785* | lncRNA | Putative role in transcription regulation | Not tissue specific (https://ngdc.cncb.ac.cn/lncbook/gene?geneid=HSALNG0140785) | n.a. |
| | | *ensembl ID ENSR00002105690 (a.k.a. ENSR00000917836) (co-localized with DMR chrX:153,046,451-153,046,767)* | | |
| *SSR4* | Signal Sequence Receptor Subunit 4 | mostly expressed in pancreas (via Human Protein Atlas) and the product is a part of a complex responsible for binding calcium to the endoplasmic reticulum membrane (via UniProt) | Tissue enhanced (pancreas) (https://www.proteinatlas.org/ENSG00000180879-SSR4/tissue) | https://www.uniprot.org/uniprot/P51571; https://www.proteinatlas.org/ENSG00000180879-SSR4/tissue |
| *SRPK3* | SRSF Protein Kinase 3 | specifically expressed in muscle (via Human Protein Atlas) and the product is required for normal muscle tissue development (via UniProt) | Tissue enhanced (skeletal muscle, tongue) (https://www.proteinatlas.org/ENSG00000184343-SRPK3/tissue) | https://www.uniprot.org/uniprot/Q9UPE1; https://www.proteinatlas.org/ENSG00000184343-SRPK3/tissue |
| *PLXNB3* | Plexin B3 | expressed specifically in brain (via Human Protein Atlas) and its product, Plexin-B3, is a receptor important in neurogenesis (via UniProt) | Tissue enriched (brain) (https://www.proteinatlas.org/ENSG00000198753-PLXNB3/tissue) | https://www.uniprot.org/uniprot/Q9ULL4; https://www.proteinatlas.org/ENSG00000198753-PLXNB3/tissue |
| *HSALNG0140793* | lncRNA | Putative role in transcription regulation | Tissue enhanced (brain, saliva secreting gland) (https://ngdc.cncb.ac.cn/lncbook/gene?geneid=HSALNG0140793) | n.a. |

[a] lncRNA = long non-coding RNA; SRP_RNA = signal recognition particle RNA; miRNA = micro RNA; piRNA = Piwi-interacting RNA;
[b] n.a. = not applicable
